# ADOLESCENT EXPOSURE TO LOW-DOSE THC DISRUPTS ENERGY BALANCE AND ADIPOSE ORGAN HOMEOSTASIS IN ADULTHOOD

**DOI:** 10.1101/2022.07.26.501615

**Authors:** Lin Lin, Kwang-Mook Jung, Johnny Le, Georgia Colleluori, Courtney Wood, Hye-Lim Lee, Francesca Palese, Erica Squire, Shiqi Su, Alexa Torrens, Yannick Fotio, Lingyi Tang, Clinton Yu, Qin Yang, Lan Huang, Nicholas DiPatrizio, Cholsoon Jang, Saverio Cinti, Daniele Piomelli

## Abstract

One of cannabis’ most iconic effects is the stimulation of hedonic high-calorie eating – the ‘munchies’ – yet habitual cannabis users are on average leaner than non-users. We asked whether this unexpected phenotype might result from lasting changes in energy balance established during adolescence, when habitual use of the drug often begins. We found that daily low-dose administration of cannabis’ intoxicating constituent, Δ^9^-tetrahydrocannabinol (THC), to adolescent mice causes an adult metabolic phenotype characterized by reduced fat mass, increased lean mass and utilization of fat as fuel, partial resistance to diet-induced obesity and dyslipidemia, and enhanced thermogenesis. Multi-omics analyses revealed that this phenotype is associated with multiple molecular anomalies in the adipose organ, which include ectopic overexpression of muscle-associated proteins and heightened anabolic processing. Thus, adolescent exposure to THC may promote an enduring ‘pseudo-lean’ state that superficially resembles healthy leanness but might in fact be rooted in adipose organ dysfunction.

## INTRODUCTION

The endogenous cannabinoid system (ECS) is found in most, if not all, mammalian organs and is involved in a diversity of physiological processes (Lu and Mackie, 2016; Piomelli and Mabou Tagne, 2022). It is comprised of two G protein-coupled receptors (CB_1_ and CB_2_), two lipid-derived ligands [anandamide and 2-arachidonoyl-*sn*-glycerol (2-AG)], and various enzymes and transporter proteins responsible for the formation, transmembrane transfer, and degradation of such ligands (Lu and Mackie, 2016; Piomelli and Mabou Tagne, 2022). All constituents of the ECS are represented in white and brown adipose tissues – the parenchymal components of the adipose organ (Cinti, 2018) – where endocannabinoid substances may act as ‘thrifty’ autocrine/paracrine messengers (DiPatrizio and Piomelli, 2012; Piazza et al., 2017) to enhance lipogenesis and attenuate non-shivering thermogenesis (Mazier et al., 2015; Ruiz de Azua and Lutz, 2019; Jung et al., 2022). The critical roles of the ECS in adipose homeostasis and energy balance are underscored by the remarkable anti-obesity effects produced, in preclinical studies and clinical trials, by globally or peripherally active CB_1_ receptor inverse agonists and neutral antagonists (Quarta and Cota, 2020). The pharmacological efficacy of these agents is primarily rooted in their ability to stimulate lipolysis, energy dissipation, and fatty acid oxidation by countering energy-conserving endocannabinoid signals in adipose, liver, and other organs (Quarta and Cota, 2020; Tam et al., 2018).

The ECS is the molecular target of the psychoactive phytocannabinoid, Δ^9^-tetrahydrocannabinol (THC) (Mackie, 2008), which accumulates in fat depots at concentrations that can fully engage local CB_1_ receptors (Kreuz and Axelrod, 1973; Torrens et al., 2020). Given the receptors’ roles in adipose function, this distribution predicts that persons who regularly use cannabis should exhibit, all else being equal, body mass index (BMI) values higher than the general population. The opposite appears, however, to be the case. Cross-sectional and longitudinal studies have consistently reported lower BMI in healthy cannabis users compared to non-users, as well as inverse associations of cannabis use with BMI, waist circumference, and other cardiometabolic risk factors (Meier et al., 2016, 2019; Beulaygue and French, 2016; Hayatbakhsh et al., 2010; Barré et al., 2021; Muniyappa et al., 2013; Clark et al., 2018; Alshaarawy and Anthony, 2019; Le Strat and Le Foll, 2011; Brook et al., 2013; Danielsson et al., 2016; Rajavashisth et al., 2012; Smit and Crespo, 2001; Penner et al., 2013; Thompson and Hay, 2015; Warren et al., 2005). The prospective National Epidemiologic Survey on Alcohol and Related Conditions (NESARC), whose sampling was devised to yield representative estimates for the adult US population, confirmed these findings (Alshaarawy and Anthony, 2019). The metabolic phenotype of habitual cannabis users is all the more paradoxical because one of the drug’s most iconic effects is the stimulation of hedonic high-calorie food intake (the ‘munchies’) (Greenberg et al., 1976; Foltin et al., 1986; Haney et al., 2007), which does not undergo noticeable tolerance with continued use of the drug (Muniyappa et al., 2013; Roberts et al., 2019).

Regular cannabis consumption often starts in adolescence. For example, in a cohort of adult individuals who successfully recovered from cannabis use disorder (Connor et al., 2021), drug use began between 15 and 17 years of age (Kelly et al., 2017). This lifespan use pattern suggests that the metabolic phenotype of habitual adult users might reflect, at least in part, enduring molecular changes initiated by CB_1_ receptor overactivation in teenage years. Consistent with this idea – and offering new insights into the contribution of the ECS to adipose homeostasis – we report now that daily exposure to a low dose of THC during adolescence causes, in mice, a lasting metabolic state characterized by lower fat mass, higher lean mass, increased utilization of fat as fuel, partial resistance to diet-induced obesity and dyslipidemia, and impaired thermoregulation. Multi-omics analyses of the adipose organ show that this state is associated with a broad transcriptional dysregulation that results, among other changes, in a striking ectopic expression of proteins normally found in skeletal and heart muscle, and enhanced anabolic processing.

## RESULTS

### Adolescent THC exposure reduces body weight gain *via* activation of CB_1_ receptors in adipocytes

Figures 1A and 1B illustrate the body weight trajectory of adolescent male and female mice (postnatal day, PND, 30-43) treated once daily with a dose of THC (5 mg/kg, intraperitoneal, IP) that, in mice of this age, produces only a modest decrease in body temperature (∼1°C) and no change in motor activity or food intake (Torrens et al., 2020; Lee et al., 2022). THC-exposed mice of both sexes gained significantly less weight than did control animals treated only with vehicle. This effect could not be attributed to changes in growth rate (Fig. 1C, Supplemental Fig. S1A, B), locomotor activity (Fig. 1D, Supplemental Fig. S1C), feeding behavior (Fig. 1E, Supplemental Fig. S1D) or nutrient absorption (Fig. 1F), none of which was affected by the drug. Moreover, there were no statistically detectable differences in intestinal microbiome composition between the two groups (Fig. 1G, Supplemental Table S1).

**Figure 1.**
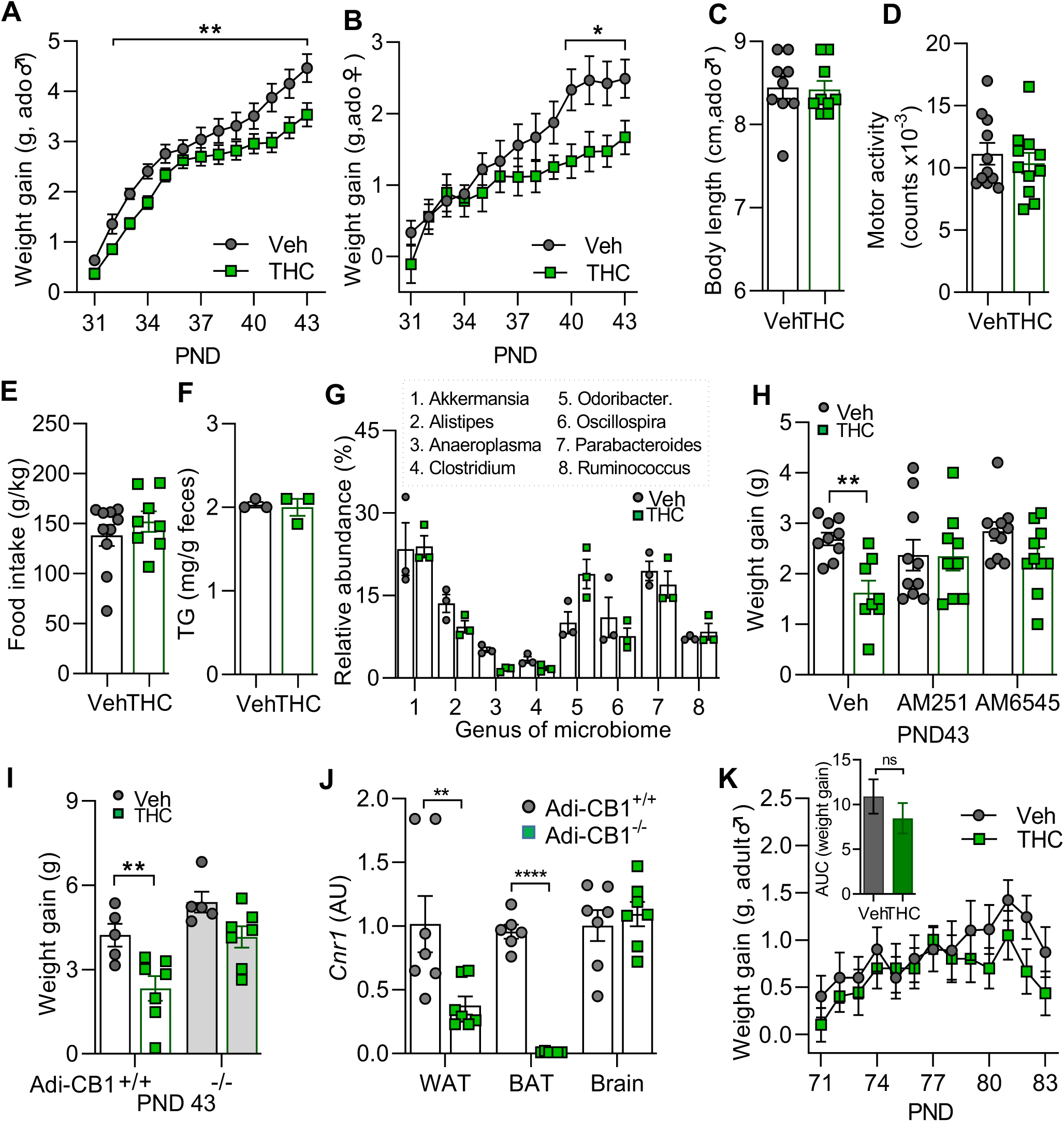
Adolescent THC exposure attenuates body weight gain via activation of CB_1_ receptors in adipocytes. **(A, B)** Effects of adolescent administration of THC (green squares) or vehicle (gray circles) on body weight gain in **(A)** male and **(B)** female mice (n=30 and 9 per group, respectively). The treatment did not affect **(C)** growth rate (n=9); **(D)** motor activity (n=11); **(E)** cumulative food intake (n=8-10); **(F)** nutrient absorption, expressed as mg of total triglycerides (TG) per g of feces (n=3 cages of 4 mice each); or **(G)** intestinal microbiome composition (n=3 cages). Displayed are bacterial genera that represent >1% of the total microbiome community. **(H)** The global CB_1_ inverse agonist AM251 (1 mg/kg) or the peripheral CB_1_ neutral antagonist AM6545 (3 mg/kg) prevented THC’s effect on body weight gain, assessed on PND43 (n=8-10). **(I)** Adolescent THC treatment did not alter weight gain in adipocyte-selective *Cnr1^-/-^* (Adi-CB_1_ ^-/-^) mice. **(J)** *Cnr1* mRNA levels in white adipose tissue (WAT), brown adipose tissue (BAT), and brain of Adi-CB_1_ ^-/-^ and Adi-CB ^+/+^ mice (n=5-7). **(K)** Subchronic THC treatment did not affect weight gain in young adult male mice (PND70-83) (n=9-10). AUC, area under the curve; ns, not significant. *P < 0.05, **P < 0.01, and ****P < 0.0001, two-way ANOVA followed by Tukey post hoc test (A,B), Student’s *t* test (C-F,J, and inset in K), mixed-effects ANOVA followed by Bonferroni’s post hoc test (H,K), or mixed-effects ANOVA followed by Bonferroni’s post hoc test **(I)**. Statistics for microbiome analyses are described under STAR Methods. See also Supplemental Figures S1, S2 and Table S1.

Three datasets indicate that adolescent THC exposure dampened weight gain through activation of CB_1_ receptors in the adipose organ, a major site of THC accumulation (Kreuz and Axelrod, 1973; Torrens et al., 2020). First, the effect of THC was prevented by co-administration of either the global CB_1_ inverse agonist AM251 (1 mg/kg, IP) or the peripherally restricted CB_1_ neutral antagonist AM6545 (3 mg/kg, IP) (Fig. 1H). At the selected dosages, neither agent affected weight gain when administered alone. Second, the response to THC was not prevented by the CB_2_ inverse agonist AM630 (1 mg/kg, IP) (Supplemental Fig. S2). Third, adolescent THC treatment did not reduce weight gain in conditional knockout mice in which the CB_1_ receptor was deleted postnatally in white and brown adipocytes (Fig. 1I). The mutants were generated by crossing CB_1_-floxed mice with transgenic AdipoqCreERT2 mice – which express Cre recombinase under the control of the promoter for the differentiated adipocyte gene *Adipoq* (Hu et al., 1996; Emont et al., 2022) – followed by tamoxifen-induced recombination on PND14-18 (Ruiz de Azua et al., 2017). Adipose-specific CB_1_ deletion was confirmed by RT-PCR (Fig. 1J). Lastly, but importantly, the same THC regimen that blunted weight gain during adolescence had no such effect when carried out in young adulthood (PND70-PND83) (Fig. 1K). The results indicate that daily exposure to low dose THC attenuates body weight gain in adolescent mice through a developmentally regulated mechanism that depends on CB_1_ receptor activation in differentiated adipocytes.

### Adolescent THC exposure modifies energy metabolism and body mass composition in adulthood

Consistent with the observed reduction in weight accrual, metabolic studies conducted at the end of the treatment period (PND44) showed that male THC-exposed mice exhibited, compared to vehicle controls, heightened energy expenditure (EE) and reduced respiratory exchange ratio (RER) during both dark (active) and light (inactive) phases of the 24-h cycle (Fig. 2A, B). Magnetic resonance imaging (MRI) analyses did not detect significant differences in fat mass or lean mass between the two groups (Fig. 2C). Total and free water content was also unchanged (Supplemental Fig. S1E1). The body weight of THC-treated animals returned to normal within approximately 10 days of treatment termination (Fig. 2D).

**Figure 2.**
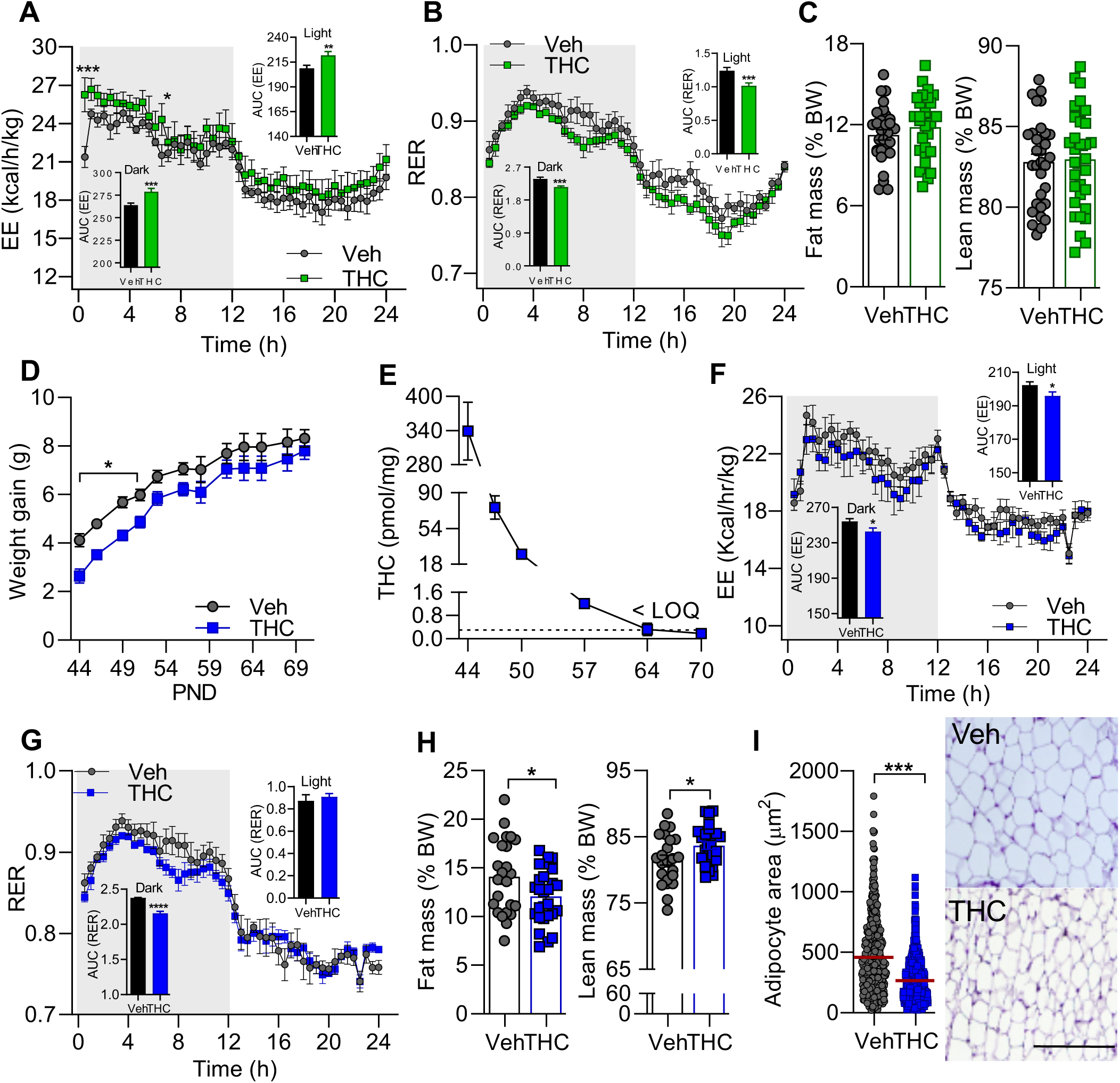
Adolescent THC exposure modifies energy metabolism and body mass composition in adulthood. **(A-C)** Residual effects of adolescent administration of THC (green squares) or vehicle (gray circles) on **(A)** energy expenditure (EE, n=4 per group); **(B)** respiratory exchange ratio (RER) (n=4); and **(C)** percent fat (left) and lean (right) mass (n=30). Measurements were made at PND44. AUC, area under the curve. **(D)** Body weight trajectory after THC treatment termination (n=15-43). **(E)** Time-course of THC concentration in epidydimal WAT following treatment termination (n=3-4); Dotted line indicates limit of quantification (LOQ). **(F-I)** Persistent effects of adolescent administration of THC or vehicle on **(F)** EE (n=4); **(G)** RER (n=4), **(H)** percent fat (left) and lean (right) mass (n=24-26); and **(I)** white adipocyte area (epididymal fat) [n=557 and 994 cells for Veh and THC, respectively, measured in three randomly selected regions (200 x 200 µm) from each mouse (n=3 mice per group)]. Bar: 100 µm. PND70 mice were used in these experiments. *P < 0.05, **P < 0.01, ***P < 0.001, and ****P < 0.0001, two-way ANOVA followed by Tukey post hoc test (A,B,F,G), Student’s *t* test (C,H,I, and insets in A,B,F,G), or mixed-effects ANOVA followed by Bonferroni’s post hoc test (D,E). See also Supplemental Figures S1, S3 and Tables S2, S3.

By young adulthood (PND70), when adipose THC content had fallen below the quantification limit of our mass spectrometry assay (0.6 pmol) (Fig. 2E), a new set of changes in energy metabolism and body-mass composition had emerged. Both EE and RER were attenuated in THC-treated mice – the former throughout the 24-h period (Fig. 2F) and the latter only during the dark phase (Fig. 2G). In addition, percent fat mass and adipocyte area in epidydimal white adipose tissue (WAT) were decreased, whereas percent lean mass was increased in the THC group compared to control (Fig. 2H-I). Total and free water content was not affected (Supplemental Fig. S1E2). Similarly, microbiome composition (Supplemental Fig. S1F) and blood chemistry profile (Supplemental Table S2) were not changed. The data suggest that adolescent exposure to low-dose THC caused in male mice two temporally distinct sets of metabolic modifications (summarized in Supplemental Table S3). At PND44, when the adipose organ still contained bioactive concentrations of the drug (∼120 nM in WAT, Fig. 2E), body weight was blunted, EE was elevated, and RER was reduced. After THC had been completely eliminated and the animals had reached young adulthood, a different metabolic state took effect in which average body weight returned to normative values, while fat mass, white adipocyte area, EE and RER were decreased, and lean mass was increased.

To better understand the lasting consequences of adolescent THC exposure on energy balance, we treated adolescent male mice with the drug or its vehicle and, when the animals had reached PND57, we gave them access to a high-fat diet (HFD, 60% kcal fat) for two months. The results show that THC-exposed animals gained significantly less weight than did vehicle controls (Fig. 3A) even though the two groups did not differ in food intake, motor activity, or nutrient absorption (Fig. 3B-D). Metabolic analyses revealed that HFD-fed mice that had received THC exhibited, compared to controls, (i) higher dark-phase EE and lower light-phase RER (Fig. 3E, F); (ii) lower percent fat mass and white adipocyte area (Fig. 3G, H); (iii) higher percent lean mass (Fig. 3I); (iv) lower fasted plasma triglycerides, cholesterol, and glucose (Fig. 3J-L); and (v) no change in water content (Supplemental Fig. S1E3; phenotype summarized in Supplemental Table S3). Glucose handling did not differ between the two groups (Fig. 3M). In separate cohorts of animals, we assessed the acute response to decreased ambient temperature, in which brown and beige adipocytes play a pivotal role (Chouchani et al., 2019). Under baseline conditions, core body temperature was ∼1°C higher in THC-exposed than control animals (Fig. 4A). After transfer to a cold room, body temperature decreased in both groups and plateaued, 2 h later, at a significantly higher level in THC-treated than control mice (Fig. 4B). Locomotor activity in the novel environment of the cold room did not differ between the two groups (Fig. 4C). Collectively, the results indicate that adolescent treatment with low-dose THC produces in male mice a lasting metabolic state characterized by lower fat mass, higher lean mass, partial resistance to diet-induced obesity and dyslipidemia, and heightened thermogenesis.

**Figure 3.**
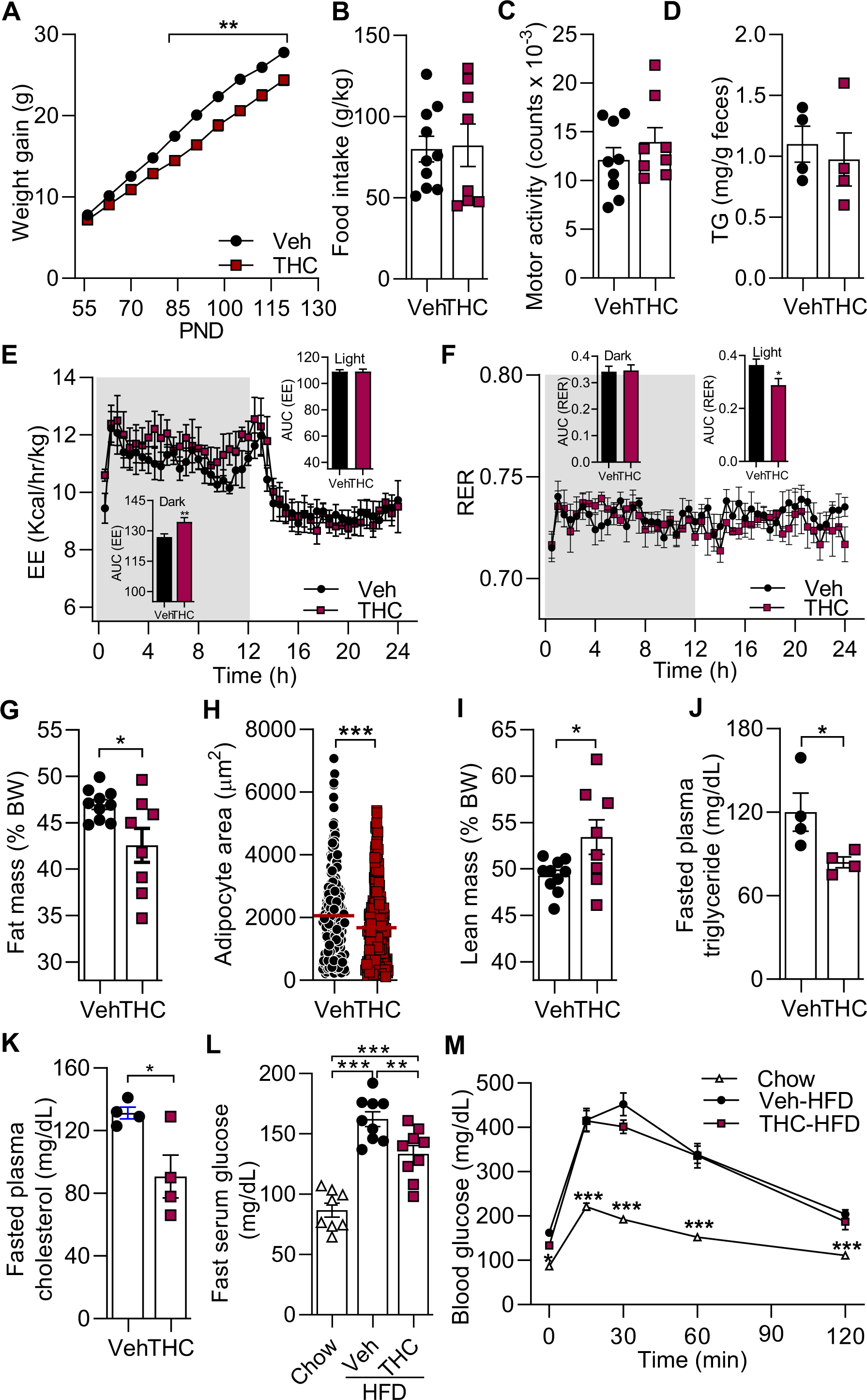
Adolescent THC exposure attenuates body-weight gain and dysmetabolic responses in adult HFD-fed mice. Persistent effects of adolescent administration of THC (red squares) or vehicle (filled circles) on metabolic responses to a HFD. **(A)** Body weight gain (n=8-11 per group); **(B)** cumulative food intake (n=8-10); **(C)** motor activity (n=8-9); **(D)** nutrient absorption expressed as mg of total triglycerides (TG) per g of feces (n=4 cages with 4 mice each); **(E)** EE (n=4); **(F)** RER (n=4); **(G)** percent fat mass (n=8-10); **(H)** white adipocyte area (epididymal fat) [n=275 and 341 cells for Veh and THC, respectively, measured in six randomly selected regions (200 x 200 µm) from each mouse (n=4 mice per group)]; **(I)** percent lean mass (n=8-10); **(J)** fasting plasma triglycerides (n=4); **(K)** fasting plasma total cholesterol (n=4); **(L)** fasting serum glucose (n=8-9); **(M)** glucose tolerance test (n=8-9). AUC, area under the curve. PND130 mice were used in these experiments. *P<0.05, **P<0.01, ***P<0.001, mixed-effects ANOVA followed by Bonferroni’s post hoc test (A,L,M), Student’s *t* test (B-D, G-K, and insets in E,F), or two-way ANOVA followed by Tukey post hoc test (E, F). See also Supplemental Figure S1 and Table S3.

**Figure 4.**
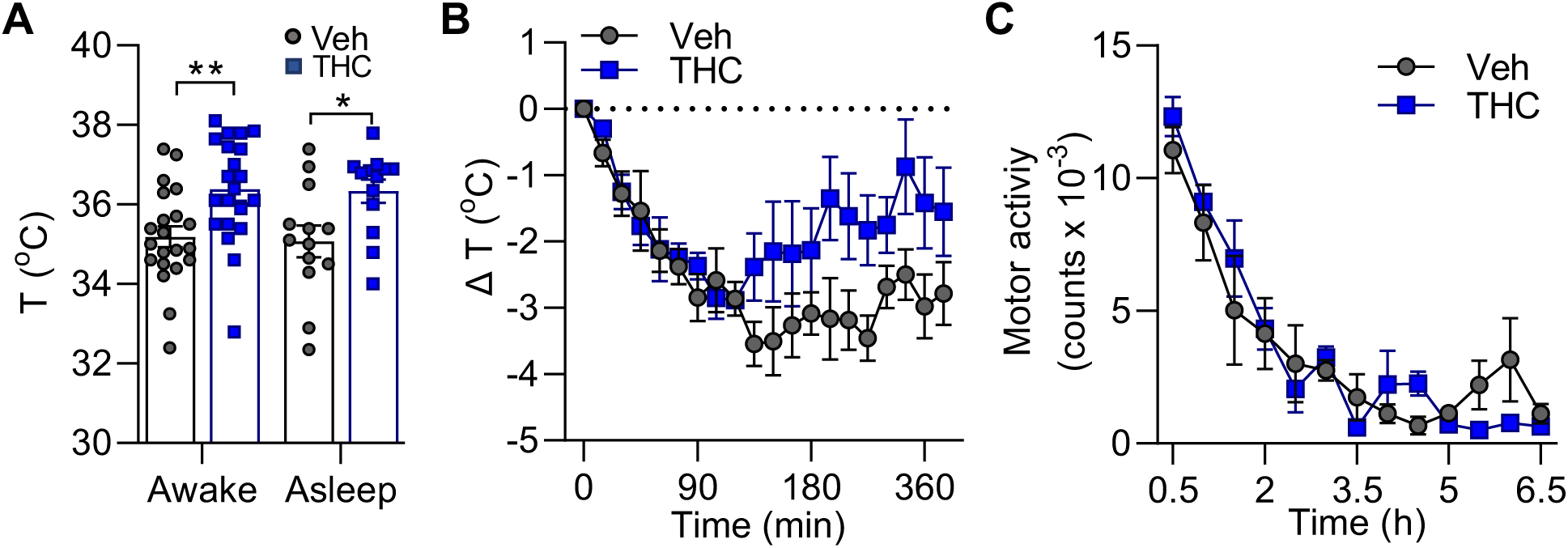
Adolescent THC exposure alters thermoregulation in adulthood. **(A)** Persistent effects of adolescent administration of THC (blue squares) or vehicle (gray circles) on core body temperature (T) in awake (n=21 per group) or asleep (n=13) adult mice housed at room temperature. Time-course of **(B)** body temperature change (ΔT) and **(C)** motor activity in THC- and vehicle-treated mice placed in a cold room for 6.5 h (n=4). Experiments were conducted on PND70. *P < 0.05, and **P < 0.01, Student’s *t* test (A), mixed-effects ANOVA followed by Bonferroni’s post hoc test (B), or two-way ANOVA followed by Bonferroni’s post hoc test (C).

### Effects of adolescent THC exposure on endocannabinoid signaling in the adult adipose organ

We next asked whether CB_1_ receptor downregulation in white and brown adipocytes, which is expected to decrease lipogenesis and stimulate thermogenesis (Mazier et al., 2015; Ruiz de Azua and Lutz, 2019; Jung et al., 2022; Clark et al., 2018), might underpin the persistent metabolic state induced by THC. Countering this possibility, however, analyses of brown adipose tissue (BAT) and WAT from THC-treated and control mice at PND70 revealed no difference in the expression of critical ECS components, including CB_1_ receptor mRNA and protein (Supplemental Fig. S3A, B) as well as mRNAs encoding for 2-AG-metabolizing enzymes, diacylglycerol lipase-α (DGL-α) and monoglyceride lipase (MGL) (Supplemental Fig. S3C, D). Accordingly, 2-AG levels were unaffected in the two tissues (Supplemental Fig. S3G, H). Transcription of N-acyl-phosphatidyl-ethanolamine phospholipase D (NAPE-PLD) and fatty acid amide hydrolase (FAAH) – which produce and deactivate, respectively, anandamide and other non-cannabinoid lipid amides (Lu and Mackie, 2016; Piomelli and Mabou Tagne, 2022) – was increased in WAT, but not in BAT (Supplemental Fig. S3E, F). Significant, but small, changes were observed in the levels of anandamide (up in WAT, down in BAT) (Supplemental Fig. S3G, H). Overall, the limited impact of adolescent THC administration on endocannabinoid signaling is unlikely to account for the treatment’s long-term effects on energy metabolism.

### Adolescent THC exposure disrupts gene transcription in the adult adipose organ

RNA sequencing experiments provided unexpected insights into the molecular events that might underpin such effects. Transcriptome datasets from interscapular BAT and epidydimal WAT of vehicle and THC-treated male mice were readily distinguishable by principal component analysis (Fig. 5A, B and Supplemental Table S4). The total number of genes differentially expressed between the two groups was 2985 (1443 up) in BAT and 3446 (1617 up) in WAT (Fig. 5D, E). Gene ontology (GO) annotation revealed, in both compartments, a striking overrepresentation of genes encoding for contractile proteins that are normally expressed in striated muscle (Fig. 5G, H). GO categories most affected were contractile fiber [GO:0043292; adjusted P value (Padj)=3.16e-49 and 1.41e-15, for BAT and WAT, respectively], myofibril (GO:0030016, Padj=3.13e-48 and 1.41e-15), sarcomere (GO:0030017, Padj=6.56e-48 and 3.43e-15), and contractile fiber part (GO:0044449, Padj=3.16e-49 and 7.67e-15). Conversely, genes involved in mitochondrial respiration and protein synthesis were underrepresented, including mitochondrial protein complex (GO:0098798, Padj=1.54e-30 and 1.19e-56, for BAT and WAT, respectively), mitochondrial inner membrane (GO:0005743, Padj=2.62e-33 and 9.89e-47), and mitochondrial matrix (GO:0005759, Padj=1.88e-19 and 3.77e-43) (Fig. 5G, H). Little or no change was seen in gene families involved in macrophage activation, innate immunity, and inflammation (Supplemental Table S4), which may be recruited when adult rats are repeatedly exposed to THC (Wong et al., 2012). The effect of THC was organ-specific because parallel analyses of skeletal muscle (hind limb quadriceps) revealed a more limited (Fig. 5C, F) and distinct (Fig. 5I) set of transcriptional alterations, which included downregulation of many of the muscle-associated genes that were elevated in BAT and WAT.

**Figure 5.**
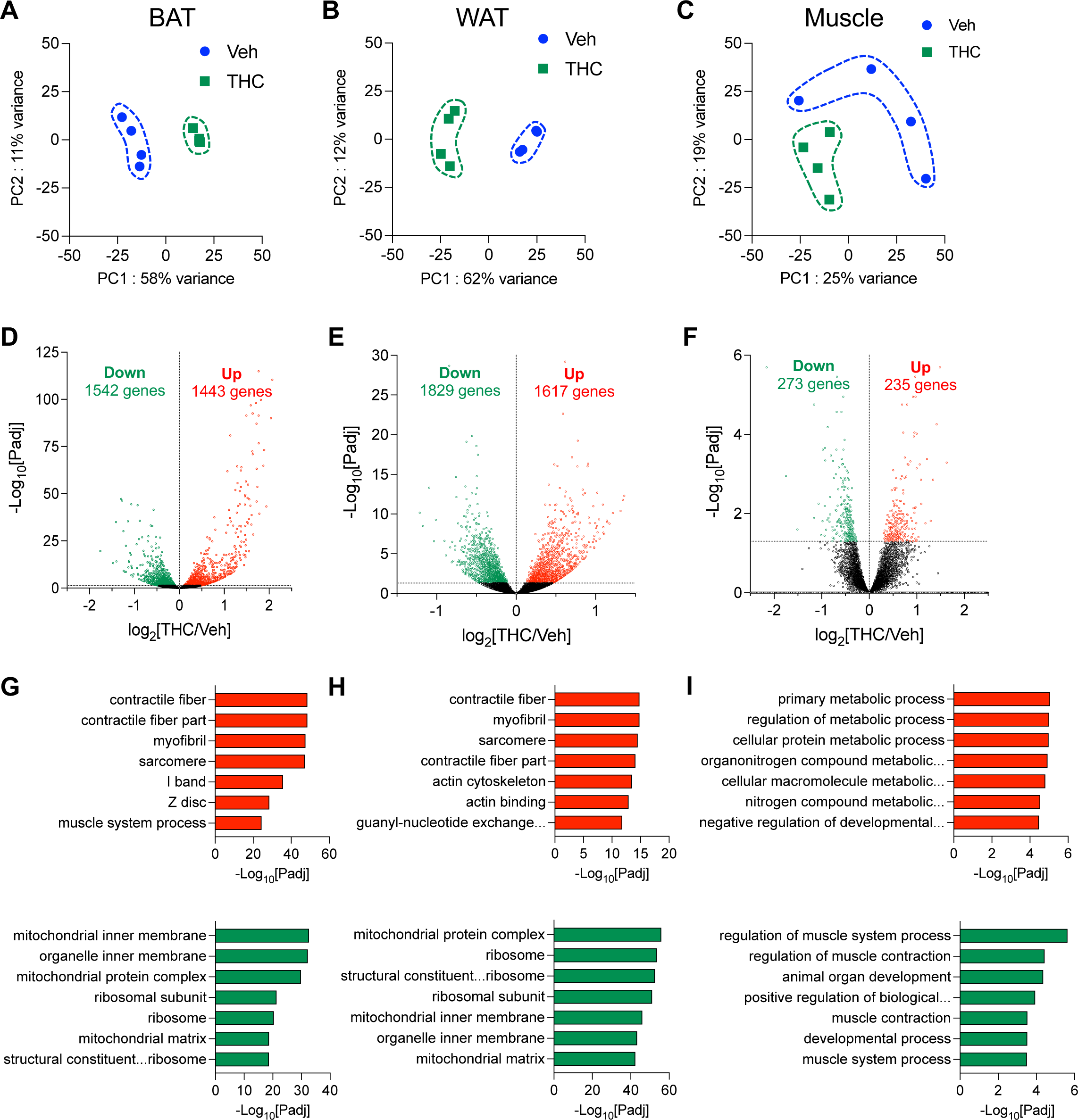
Adolescent THC exposure disrupts gene transcription in adult adipose organ. **(A-C)** Principal component analysis of transcriptome datasets from **(A)** BAT, **(B)** WAT, and **(C)** skeletal muscle of PDN70 mice treated in adolescence with THC (green squares) or vehicle (blue circles). **(D-F)** Volcano plots showing genes differentially expressed (P_adj_<0.05) in **(D)** BAT, **(E)** WAT, and **(F)** skeletal muscle. Red, upregulated; green, downregulated; black, unchanged (P_adj_>0.05). **(G-I)** GO categories showing highest enrichment in **(G)** BAT, **(H)**, WAT, and **(I)** skeletal muscle. Ranking is according to –log_10_ P_adj_ value. Top, upregulated genes (red bars); bottom, downregulated genes (green bars). N=3-4 per group. Statistical analyses are described under STAR Methods. See also Supplemental Table S4.

Closer inspection of the data showed that transcription of multiple genes encoding for sarcomere constituents – e.g., titin (*Ttn*), myosin heavy chain (*Myh*) 1, 2 and 7, myosin light chain 3 (*Myl2*), troponin 1 (*Tnni2*), and troponin T (*Tnnt3*) – as well as enzymes primarily found in muscle – e.g., enolase 3 (*Eno3*) and sarco(endo)plasmic reticulum calcium ATPase 1(*Atp2a1*) – was markedly increased in BAT and, to a lesser extent, in WAT (Fig. 6A). Conversely, transcription of nuclear-encoded genes involved in mitochondrial respiration – including *Atp5* (ATP synthase subunit 5) and *Ndufa* (NADH:ubiquinone oxidoreductase), two components of respiratory chain complex I – was suppressed (Fig. 6B). It is worth noting, however, that expression of the master regulator of mitochondrial biogenesis, *Pgc1a* (peroxisome proliferator-activated receptor-γ coactivator-1α), was enhanced in both BAT and WAT (Supplemental Fig. S4A), while transcription of multiple genes that are essential for adipose organ function was either unaffected (peroxisome proliferator-activated receptor-γ, *Pparg*) or differentially regulated in the two tissues [PR/SET Domain 16 (*Prdm16*), Δ-adrenergic receptor (*Adrb3*), uncoupling protein 2 (*Ucp2*): down in BAT, up in WAT; *Ucp1*: up in BAT, unchanged in WAT; *Ucp3*: unchanged in BAT, up in WAT] (Supplemental Fig. S4B-G). Confirming the organ-specificity of THC’s effects, a distinct and partially opposite set of transcriptional modifications was evident in skeletal muscle, where many genes implicated in sarcomere structure/function, calcium transport, and cellular respiration were down-regulated in THC-exposed relative to control animals (Fig. 6A, B). Thus, adolescent THC treatment produces a complex array of enduring transcriptional abnormalities in the mouse adipose organ – including ectopic overexpression of genes encoding for muscle proteins, and suppression of genes involved in mitochondrial respiration.

**Figure 6.**
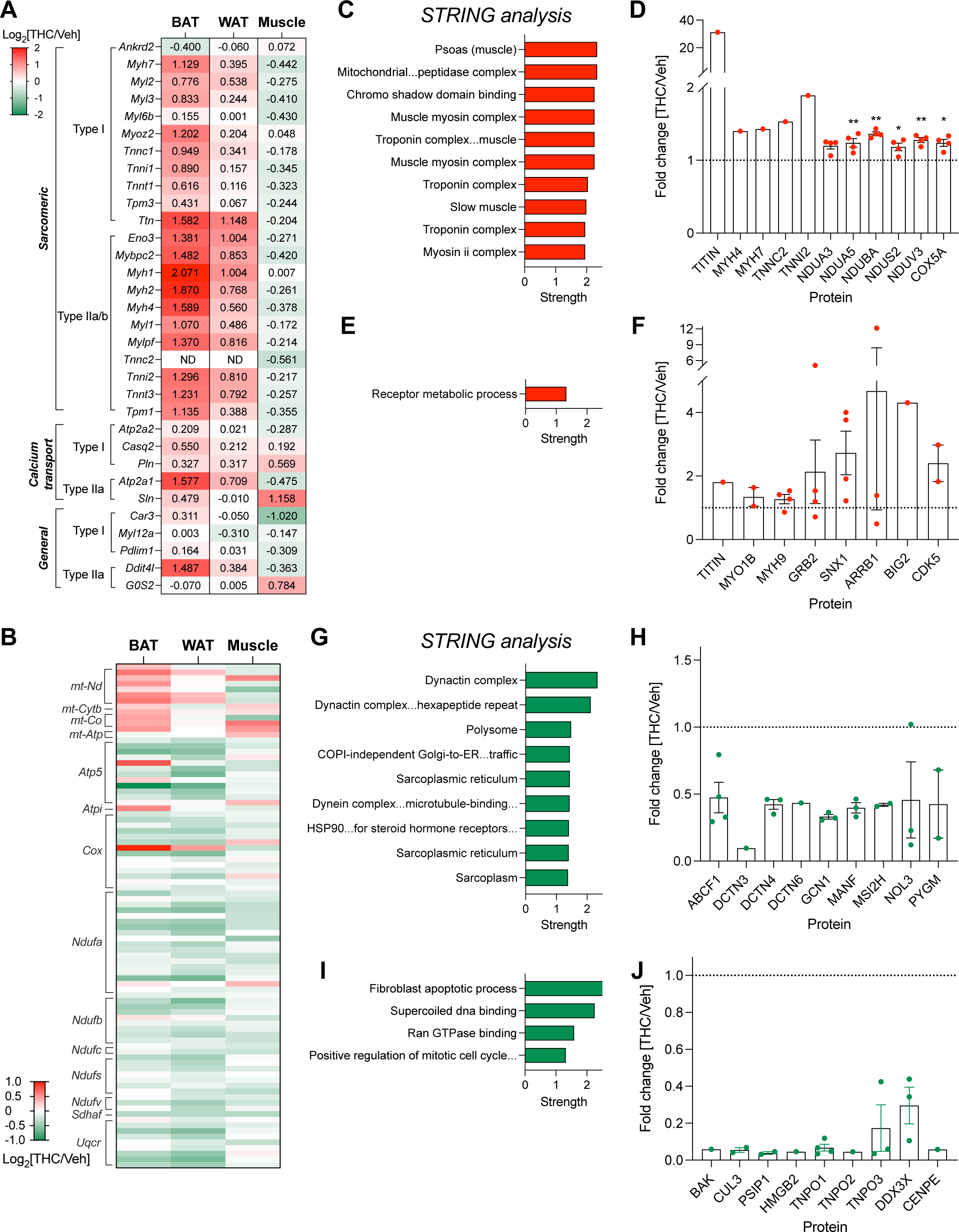
Adolescent THC exposure causes ectopic expression of muscle-associated proteins in adult adipose organ. **(A-B)** Heatmaps showing the effect of adolescent THC treatment on the transcription of select **(A)** muscle-associated genes, and **(B)** genes involved in mitochondrial function and oxidative phosphorylation. In (A), numbers represent the log_2_[THC/Veh] value for each gene. **(C-J)** Untargeted proteomics analyses for BAT (C, D, G, H) and WAT (E, F, I, J). STRING annotation of proteins (**C, E**) upregulated (red) or (**G, I**) downregulated (green) in THC- vs vehicle-treated mice at PND70. Protein term IDs showing significant enrichment (P_adj_<0.05) were identified and those with strength >1.0 are ranked in figure. **(D, F, H, J)** Fold changes in abundance of select proteins that are **(D, F)** upregulated (red) or **(H, J)** downregulated (green) by THC treatment. *P < 0.05, and **P < 0.01 by student’s *t* test (n=4) (D,F,H,J). See also Supplemental Figures S4, S5 and Tables S5-S7.

### Adolescent THC exposure alters protein expression in the adult adipose organ

Consistent with the transcriptomic data, untargeted mass spectrometry analyses revealed substantial proteomic alterations in adipose organ of adult male mice that had received THC in adolescence. Statistically detectable differences (*P* < 0.05) between THC-exposed and control animals were observed in both BAT (102 proteins, 75 up) and WAT (44 proteins, 6 up) (Supplemental Fig. S5A, B). In BAT of THC-treated mice, STRING analysis (Szklarczyk et al., 2021) revealed, among proteins with high fold-increases, a marked enrichment (strength > 1.9) in categories related to skeletal muscle components (Fig. 6C, Supplemental Table S5). Affected proteins included essential sarcomere constituents such as titin (∼30-fold increase), troponin (∼2- fold), and myosin (∼1.5-fold) (Fig. 6D) – whose gene transcription was enhanced (Fig. 6A) – but also proteins involved in mitochondrial respiration, such as NADH dehydrogenase 1 α- subcomplex subunit 3 and 5 (NDUA3 and 5), and cytochrome C oxidase subunit 5A (COX5A) (Fig. 6D) – whose transcription was suppressed (Fig. 6B). Additional proteins overrepresented in BAT of THC- treated mice are listed in Supplemental Figure S5C. Proteins exclusively detected in THC-exposed BAT, 160 in total, are reported in Supplemental Table S6, while down-regulated proteins are listed in Figure 6G, H and Supplemental Table S7. THC treatment produced fewer proteomic changes in WAT than it did in BAT (Supplemental Fig. S5B). Nevertheless, the levels of three muscle-associated proteins – titin, myosin 1B, and myosin 9 – were noticeably increased in this tissue (Fig. 6E, F). Other proteins overrepresented in WAT of THC-treated mice are reported in Figure 6F and Supplemental Table S5. Moreover, proteins only detectable in THC-exposed WAT, a total of 288, are shown in Supplemental Table S6. Downregulated proteins are listed in Figure 6I, J and Supplemental Table S7.

### Adolescent THC exposure modifies amino acid metabolism in adult BAT

Targeted mass spectrometry analyses revealed significant upregulation of several metabolites in BAT of THC-exposed mice (Fig. 7A-C). Unexpectedly, metabolites related to energy production pathways such as glycolysis, tricarboxylic acid cycle, and pentose phosphate pathway were not affected by THC treatment (Supplemental Fig. S6A-C). Nucleotides were also largely unchanged (Supplemental Fig. S6D). By contrast, essential and non-essential amino acids, as well as N-acetylated amino acids, were elevated in the THC group (Fig. 7D-F), which is suggestive of enhanced anabolic processing. Consistent with this idea, we found that NADPH/NADP^+^ ratio was significantly decreased (Fig. 7G), possibly due to increased NADPH consumption for protein synthesis. The NADH/NAD^+^ ratio was unchanged (Fig. 7G). Interestingly, no significant metabolomic alterations were seen in WAT (Supplemental Fig. S6E-K), in which the transcriptomic and proteomic response to THC was also less pronounced. Finally, observations by transmission electron microscopy found no significant ultrastructural difference in brown and white adipocytes from THC-exposed and control mice (Supplemental Figs. S7A-C). Furthermore, in BAT, no differences were seen in the density of parenchymal noradrenergic (tyrosine hydroxylase-positive) fibers or lipid droplet size (Supplemental Fig. S7D).

**Figure 7.**
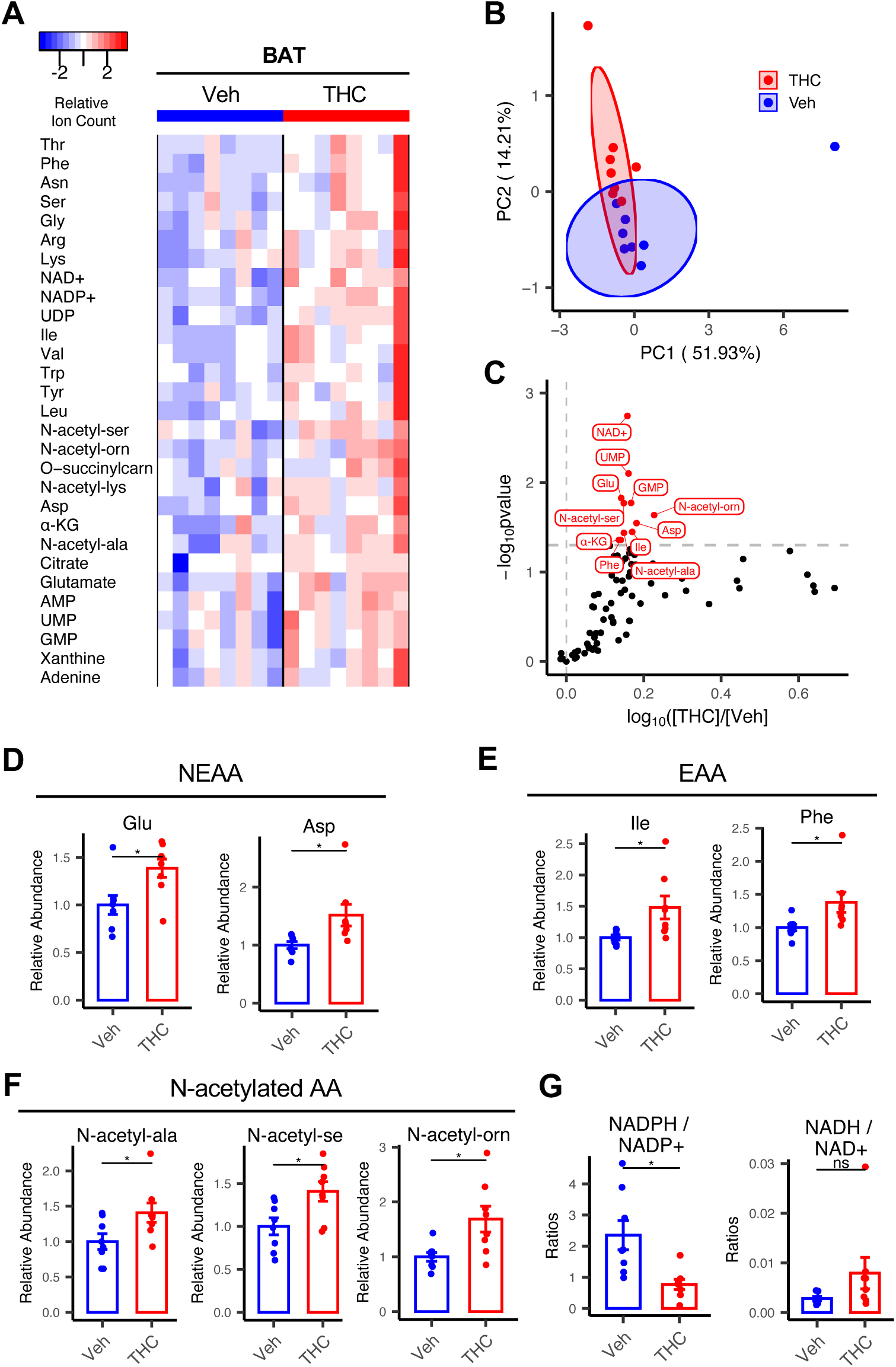
Adolescent THC exposure modifies amino acid metabolism in adult BAT. **(A)** Heatmap illustrating the effects of adolescent THC treatment on select intermediate metabolites in BAT. Top 29 compounds with highest statistical significance are shown (Student’s *t* test, n=8 per group). **(B)** Principal component analysis of top 29 compounds. **(C)** Volcano plot showing changes in individual metabolites (log_10_ of THC/Veh). **(D-F)** Relative abundance of the indicated metabolites. **(G)** NADPH/NADP^+^ and NADH/NAD^+^ ratios. *P < 0.05 by Student’s *t* test (n=8). See also Supplemental Figure S6.

## DISCUSSION

In this study, we examined whether adolescent exposure to the psychotropic constituent of cannabis, THC, alters energy balance and adipose organ function in adult life. The question is important for at least two reasons. First, a significant fraction of teenagers use cannabis regularly. In 2021, prevalence levels of daily cannabis consumption in US schools were 0.6%, 3.2%, and 5.8%, in 8^th^, 10^th^, and 12^th^ grade, respectively (Miech et al., 2022). Second, epidemiological studies have demonstrated a robust association between habitual cannabis use and alterations in metabolic function (Meier et al., 2019; Beulaygue and French, 2016; Hayatbakhsh et al., 2010; Barré et al., 2021; Muniyappa et al., 2013; Clark et al., 2018; Alshaarawy and Anthony, 2019; Le Strat and Le Foll, 2011; Brook et al., 2013; Danielsson et al., 2016; Rajavashisth et al., 2012; Smit and Crespo, 2001; Penner et al., 2013; Thompson and Hay, 2015; Warren et al., 2005). Understanding the molecular bases of such alterations and their possible link to adolescent exposure is necessary to inform evidence-based prevention and guide future research.

We found that a daily low-intensity THC regimen (Torrens et al., 2020; Lee et al., 2022) dampened body weight gain in adolescent but not young adult mice. When adolescent THC treatment was stopped, the animals weighed less and expended more energy than did vehicle-treated controls. By the time they reached adulthood, mice in the THC group transitioned to a different metabolic state whose attributes included normal body weight, decreased fat mass and white adipocyte area, increased lean mass, partial resistance to diet-induced obesity and dyslipidemia, and heightened thermogenesis. Utilization of fat as fuel was increased during the dark (active) phase of the 24-h cycle, but energy expenditure could be either lower or higher than control, depending on whether the mice were fed a normal chow or a high-fat diet. This state was accompanied by an aberrant omics profile characterized, in both BAT and WAT, by overexpression of genes and proteins that are normally restricted to skeletal and heart muscle and, in BAT only, by heightened anabolic processing. The results thus suggest that frequent exposure to low-dose THC during adolescence promotes a maladaptive ‘pseudo-lean’ state – that is, a state that only superficially resembles healthy leanness – which may result, at least partly, from lasting modifications of the adipose organ.

Our results suggest that THC impairs normal body weight accrual in adolescent mice through activation of CB_1_ receptors in differentiated adipocytes. Supporting this conclusion, we found that the response (i) was prevented by a globally active CB_1_ inverse agonist (AM251) or a peripherally restricted CB_1_ neutral antagonist (AM6545); (ii) was absent in mice selectively lacking CB_1_ in adipocytes that express the *AdipoQ* gene (Hu et al., 1996) and are likely, therefore, to be differentiated (Emont et al., 2022); and (iii) was not affected by the CB_2_ inverse agonist AM630. In addition to delineating the receptor mechanism through which THC affects weight gain in adolescence, these results reveal an unexpected contribution of CB_1_ receptors to adipose organ homeostasis, which appears to be distinct from their known roles in energy conservation (Mazier et al., 2015; Ruiz de Azua and Lutz, 2019; Jung et al., 2022). It is important to point out, however, that our findings do not rule out the possibility that CB_1_ receptors in organs other than the adipose might contribute, either directly or indirectly, to specific aspects of the metabolic phenotype induced by THC. In fact, we found that transcriptional programs in skeletal muscle are also significantly impacted by adolescent THC treatment, as convincingly documented for the brain (Bara et al., 2021).

Along with a pseudo-lean phenotype, adolescent THC exposure also causes pervasive transcriptional abnormalities in BAT and WAT. Many muscle-associated genes were induced in these tissues, whereas genes encoding for components of the mitochondrial respiratory chain were suppressed. Moreover, *Pgc1a* levels were elevated in both BAT and WAT, while other critical regulators of adipocyte function were differentially affected – for example, *Adrb3* was down in BAT and up in WAT, while *Ucp1* was up in BAT and unchanged in WAT. Consistent with the transcriptional data, several muscle-associated proteins – including titin, myosin, and troponin – were abnormally expressed in BAT and WAT of THC-exposed mice. Interestingly, however, key proteins involved in mitochondrial respiration (e.g., NDUA3 and 5, COX5A), whose transcription was suppressed, were increased in BAT after THC treatment. Mismatches between mRNA and protein levels are not uncommon (Buccitelli and Selbach, 2020) and are thought to reflect differences in temporal factors – such as half-life time and translation rate constant– which might be widened by THC-dependent disruption of gene transcription and protein synthesis. The anomalous molecular landscape induced by adolescent THC exposure defies straightforward interpretation. It is reasonable to speculate, however, that the adipose organ might adjust to CB_1_ receptor overactivation during the second month of postnatal life, when the rate of adipocyte proliferation is high (Greenwood and Hirsch, 1974), by entering a lasting dyshomeostatic condition in which lipid storage in WAT is defective and anabolic processing and thermogenesis in BAT are enhanced.

The link between cannabis consumption, lower BMI, and improved cardiometabolic risk is well established in the epidemiological literature (Meier et al., 2019; Beulaygue and French, 2016; Hayatbakhsh et al., 2010; Barré et al., 2021; Muniyappa et al., 2013; Clark et al., 2018; Alshaarawy and Anthony, 2019; Le Strat and Le Foll, 2011; Brook et al., 2013; Danielsson et al., 2016; Rajavashisth et al., 2012; Smit and Crespo, 2001; Penner et al., 2013; Thompson and Hay, 2015; Warren et al., 2005) and cannot be ascribed to differences in lifestyle or to a tendency of lean individuals to become cannabis users (Clark et al., 2018). In particular, longitudinal studies have shown that continuous regular use of cannabis from adolescence (12-18 years) to adulthood (32-38 years) is associated with lower BMI, smaller waist-hip ratio, lower fasting triglyceride and glucose levels, and improved cardiometabolic profile (Meier et al., 2016, 2019). To reconcile these findings with cannabis’ ability to stimulate palatable high-calorie eating (Greenberg et al., 1976; Foltin et al., 1986; Haney et al., 2007), it has been proposed that habitual use of the drug might lead to a reduction in the number and/or signaling efficiency of CB_1_ receptors. This hypothesis is inconsistent, however, with available evidence showing persistent hedonic eating (Roberts et al., 2019) and increased carbohydrate and energy intake (Smit and Crespo, 2001; Muniyappa et al., 2013) in regular cannabis users. An intriguing possibility, suggested by our data, is that the paradoxical metabolic profile of adult cannabis users might result, at least partly, from lasting modifications in adipose organ function induced by THC during adolescence.

In conclusion, the present results show that daily administration of low-dose THC in adolescent mice results in an enduring metabolic state characterized by decreased fat mass, increased lean mass, partial resistance to diet-induced obesity and dyslipidemia, and enhanced thermogenesis. Transcriptomic, proteomic, and metabolomic analyses show that this state is accompanied by a complex set of molecular anomalies in BAT and WAT, which include overexpression of proteins that are normally found in skeletal and heart muscle. Whether such state, referred to here as pseudo-lean, might impact physical, social, and mental health is an important question for future studies.

## ACKNOWLEDGEMENTS

This work was supported by the National Institutes on Drug Abuse (Grant No. P50DA044118-01 [to DP]). The authors thank the researchers of the Genomics High-Throughput Facility (GHTF) and Microbiome Center in the University of California, Irvine for assistance with microbiome analysis. The authors wish to thank Jade Ramirez for managing the animals, Dr. Faizy Ahmed for assistance with LC-MS/MS analysis, and Drs. Bruce Blumberg and Riann Egusquiz for sharing the magnetic resonance imaging (MRI) equipment.

## AUTHOR CONTRIBUTIONS

D.P., L.L., and K.-M. J. designed the studies and L.L. performed the experiments with the assistance from H.-L.L, F.P., E.S., S.S., A.T., and Y.F. J.L. and C.J. carried out metabolomic analysis, C.Y and L.H performed proteomic analysis, and G.C. and S.C. performed morphological analysis using electron microscope. C.W. and N.D. carried out experiments with conditional CB1 knockout mice. L.T. and Q.Y. provided help with the metabolic chamber study. D.P. ideated the project, supervised it, and wrote the manuscript with assistance from L.L., K.-M.J., C.J., and S.C.

## STAR METHODS

## RESOURCE AVAILABILITY

### Lead contact

Further information and requests for resources and reagents should be directed to and will be fulfilled by the Lead Contact, Dr. Daniele Piomelli.

### Materials availability

This study did not generate new unique reagents.

## EXPERIMENTAL MODEL AND SUBJECT DETAILS

### Chemicals

We purchased AM251, AM6545, AM630, oleoylethanolamide (OEA), palmitoylethanolamide (PEA), anandamide (AEA), 2-arachidonoylglycerol (2-AG), and THC from Cayman Chemicals (Ann Arbor, MI). [^2^H_3_]-THC was from Cerilliant (Round Rock, TX). Other deuterium-containing standards were from Cayman Chemicals. Liquid chromatography (LC) solvents and all other chemicals were from Sigma Aldrich (Saint Louis, MO) or Honeywell (Muskegon, MI, USA). Formic acid was from Thermo Fisher Scientific (Waltham, MA). All solvents and chemicals were of the highest available grade.

### Animals

We obtained C57BL/6J mice of both sexes from Jackson Laboratories (Farmington, CT). Conditional adipose tissue-CB_1_-knockout (Ati-CB_1_-KO) mice were generated by crossing AdipoqCreERT2 mice (Sassmann et al., 2010) with mice containing two loxP sites flanking the open reading frame of the *Cnr1* gene (Marsicano et al., 2003; Ruiz de Azua et al., 2017). Tamoxifen treatment began at PND14 and consisted of 5 daily intraperitoneal (IP) injections of 40 mg/kg in 4 mL of corn oil. Mice were transferred back to their home cages 72 h after the final tamoxifen injection. All animals were housed in ventilated cages (4-5 per cage) with food and water available ad libitum. Age- and weight-matched animals were randomly assigned to treatment groups and were allowed to acclimate for at least one week before experiments. Housing rooms were maintained on a 12-h light/12-h dark cycle (lights on at 6:30 AM) with constant temperature (20±2°C) and relative humidity (55–60%). All procedures were approved by the Institutional Animal Care and Use Committee of the University of California, Irvine, and were carried out in strict accordance with the National Institutes of Health guidelines for the care and use of experimental animals.

### Drug administration

Drugs were freshly prepared and administered by IP injection. THC was dissolved in a vehicle of Tween 80/sterile saline (5:95, v/v) and was administered once daily on PND30-43, unless specified otherwise. The pharmacokinetic and pharmacodynamic properties of the THC dose selected for the present study (5 mg/kg) are described in detail elsewhere (Torrens et al., 2020; Lee et al, 2022). AM251, AM6545 and AM630 were dissolved in dimethyl sulfoxide (DMSO) in sterile saline (5:95, v/v) and administered by IP injection 60 min before THC.

### Tissue collection

Mice were anesthetized with isoflurane (Pivetal veterinary, Loveland, CO), and cardiac blood was collected into ethylenediaminetetraacetic acid (EDTA)-rinsed syringes and transferred into 1 ml polypropylene plastic tubes containing spray-coated potassium-EDTA. Blood was centrifuged at 1,450 x *g* at 4°C for 15 min and supernatants were transferred into polypropylene tubes. The animals were decapitated and interscapular BAT, epididymal WAT, and hindlimb quadriceps muscle were quickly removed, snap-frozen and stored at -80°C until analyses.

### High-fat diet (HFD)-induced obesity

Groups of C57BL/6 mice received either vehicle (Tween 80/sterile saline, 5:95, v/v) or THC (5 mg/kg, IP) on PND30-43. Starting from PND57, they were exposed to a high-fat diet (HFD, 60 kcal % fat, D12492; Research Diets, New Brunswick, NJ) for 10 weeks. Body weight and food intake were recorded three times per week.

### Body growth

A group of PND44 mice was lightly anesthetized with isoflurane and body, tail and head length were measured using a ruler, holding the animals vertically by the tail.

### Body composition

Body composition was measured on PND44, PND70 and PND130 (HFD-exposed group), using an EchoMRI^TM^ Whole Body Composition Analyzer (EchMRI, Houston, TX) as described (Chamorro-García et al., 2018).

### Feeding and motor activity

Feeding and motor activity were recorded using an automated system (SciPro, New York, NY), as described (Gaetani et al., 2003). Mice were habituated to test cages for 5 days prior to trials. 24-h food intake (g/weight) and motor activity were measured. We assessed the following feeding parameters: average satiety ratio [min/(g/kg)], feeding latency (min), meal size (g/kg), post meal interval (min), meal frequency (meals/h) and duration (Gaetani et al., 2003).

### Energy expenditure

Mice were individually acclimated to metabolic chambers (PhenoMaster System, TSE, Germany) for 24 h. Motor activity, O_2_ consumption, and CO_2_ production were recorded at 30-min intervals for 3 consecutive days. Body weight and food intake were measured before and after the tests.

### Blood chemistry

Plasma samples (0.1 ml) were prepared from cardiac blood obtained from mice at PND44, PND70, and PND130 (HFD-exposed group) after overnight fasting. Analyses were conducted at Antech Diagnostics (Irvine, CA).

### Efficiency of nutrient absorption

Fecal matter (∼0.5 g) was collected on PND43-44 and PND69-70 from group-housed mice (4 per cage). The samples were snap frozen in liquid N_2_, pulverized in a mortar, and transferred to 16- ml glass tubes. Sterile saline (5 ml) and chloroform/methanol (2:1, v/v; 5 ml) were added to the sample, stirred vigorously, and centrifuged at 1,000 x *g* for 15 min at 4°C. The organic phases were collected and dried under N_2_. Extracts were reconstituted in NP40 substitute assay reagent (Cayman Chemical) containing protease inhibitors. Triglycerides were quantified using a colorimetric assay kit (Cayman Chemical).

### Glucose tolerance test

On PND130, HFD-exposed mice were food deprived overnight in cages equipped with a wired bottom (to prevent coprophagia). Three h prior to the test, the animals were placed in new cages in a quiet room. During the test, body weight was recorded, and 20% glucose (1 g of glucose/kg body weight) was administered by IP injection. Tail blood was collected before and 15, 30, 60 and 120 min after injections. Glucose concentrations were measured using a commercial instrument (Aviva, ACCU-CHEK, Indianapolis, IN).

### Temperature measurements

Under light isoflurane anesthesia, we implanted and fixed with surgical glue temperature microchips (United Information Devices, Lake Villa, IL) in the peritoneum of male mice at PND68. Animals were returned to their home cages for 24 h. On PND69, we recorded baseline temperature during the light and dark phases. On PND70, the animals were transferred to a walk-in cold room (4°C) and temperature was measured at periodic intervals for the following 6.5 h.

### THC measurements

#### THC extraction

Epidydimal adipose tissue (15-20 mg) was transferred into 2-ml Precellys tubes (Bertin Instruments, France) and homogenized in ice-cold acetonitrile (0.5 ml) containing 1% formic acid and [^2^H_3_]-THC (50 pmol) as internal standard. The samples were stirred vigorously for 30 s and centrifuged at 2,800 x *g* at 4°C for 15 min. Supernatants were loaded onto Captiva Enhanced Matrix Removal (EMR) cartridges (Agilent Technologies, Santa Clara, CA) prewashed with water/acetonitrile (1:4, v/v). The extracts were eluted under positive pressure (3-5 mmHg, 1 drop/5 sec) using a Positive Pressure manifold 48 processor (PPM-48, Agilent Technologies). Tissue pellets were rinsed with water/acetonitrile (1:4, v/v; 0.2 ml), stirred for 30 s, and centrifuged again at 2,800 x *g* at 4°C for 15 min. The supernatants were transferred onto EMR cartridges, eluted, and pooled with the first eluate. The cartridges were washed again with water/acetonitrile (1:4, v/v; 0.2 ml), and pressure was increased gradually to 10 mmHg (1 drop/sec) to ensure maximal analyte recovery. Eluates were dried under N_2_ and reconstituted in methanol (0.1 ml) containing 0.1% formic acid and transferred to deactivated glass inserts (0.2 ml) placed inside amber glass vials (2 ml; Agilent Technologies).

#### LC/MS-MS analysis

LC separations were carried out using a 1200 series LC system coupled with a 6410B mass spectrometric detector (Agilent Technologies). Analytes were separated on an Eclipse XDB C18 column, 1.8-μm, 2.1 x 50 mm (Agilent Technologies). The mobile phase consisted of water containing 0.1% formic acid as solvent A and methanol containing 0.1% formic acid as solvent B. The flow rate was 0.5 ml/min. The gradient conditions were as follows: starting 75% B to 89% B in 3.0 min, changed to 95% B at 3.01 min and maintained till 4.5 min to remove any strongly retained materials from the column followed by column re-equilibration with 75 % B for 2.5 min. The total analysis time, including re-equilibrium, was 7 min. The column temperature was maintained at 40°C and the autosampler at 9°C. The injection volume was 2.0 μl. To prevent carry-over, the needle was washed in the autosampler port for 30 s before each injection using a wash solution consisting of 10% acetone in water/methanol/isopropanol/acetonitrile (1:1:1:1, v/v). The mass spectrometer was operated in the positive electrospray ionization mode, and THC/internal standard were quantified by multiple reaction monitoring (MRM) using transitions listed in Methods Table M1. Capillary voltage was set at 3,500 V. Source temperature was 300°C, and gas flow was set at 12.0 l/min. Nebulizer pressure was set at 40 psi. Collision energy and fragmentation voltage were set as reported (Torrens et al., 2020). The MassHunter software (Agilent Technologies) was used for instrument control, data acquisition, and data analysis.

### Endocannabinoid measurements

#### Tissue extraction

Frozen epidydimal WAT and interscapular BAT samples (∼40 mg) were transferred to 2 ml Precellys soft tissue vials (Bertin Instruments) and ice-cold acetone (1 ml) containing [^2^H_4_]-anandamide and [^2^H_5_]-2-AG (100 nmol each) was added. The samples were homogenized at 4°C at 6,800 rpm, 15 s per cycle for two cycles with a 20-s pause. They were then centrifuged at 830 x *g* for 15 min at 4°C and supernatants were eluted over EMR-Lipid cartridges, as described above. Eluates were dried under N_2_, reconstituted in acetonitrile (0.1 ml), and stored at -20°C until analysis.

#### LC/MS-MS analysis

Endocannabinoids were fractionated using a 1260 series LC system (Agilent Technologies) coupled to a 6460C triple-quadrupole mass spectrometric detector (MSD; Agilent), as described (Lin et al., 2022). A step gradient separation was performed on a Poroshell 120 column 1.9 µm, 2.1 x 100 mm (Agilent Technologies) with a mobile phase consisting of 0.1% formic acid in water as solvent A and 0.1% formic acid in acetonitrile as solvent B. A linear gradient was used: 0.0–9.5 min 55.0 % B - 80% B; 9.51-11.0 min 95% B; and 11.01–15.50 min maintained at 55% B for column re-equilibration. Column temperature was 40°C and autosampler temperature at 9°C. Injection volume was 2 µl, flow rate was 0.3 ml/min, and total analysis time, including column re-equilibration, was 15.5 min. To prevent carry-over, the needle was washed in the autosampler port for 30 s before each injection using a wash solution consisting of 10% acetone in water/methanol/isopropanol/acetonitrile (1:1:1:1, v/v). The mass spectrometer was operated in the positive electrospray ionization mode, and analytes were quantified using the MRM transitions listed in Methods Table M1. Capillary and nozzle voltages were set at 3,500 V and 500V respectively. Drying gas and sheath temperatures were 300°C with gas flows of 9.0 L/min and 12 L/min. Nebulizer pressure was set at 40 psi. MassHunter software (Agilent Technologies) was used for instrument control, data acquisition, and data analysis.

### Morphological analyses

#### Measurement of adipocyte area

Epididymal WAT was fixed by overnight incubation in phosphate-buffered saline (PBS, 0.1 M, pH 7.4) containing 4% paraformaldehyde (PFA). The samples were rinsed, dehydrated, cleared, and embedded in paraffin. Sections (7 μm thickness) were prepared using a Leica microtome RM2255 (Leica Biosystems, Deer Park, IL). Sections were dehydrated in ethanol, cleared, mounted, and stained with hematoxylin/eosin (National Diagnostics, Atlanta, GA). Transilluminated images of H&E-stained tissues were collected using a Ti Eclipse Microscope (Nikon, Melville, NY) with a Plan Apo 10 x objective. Images were analyzed using the NIH image J software.

#### Immunohistochemistry

Samples were processed as described (Colleluori et al., 2022) . Briefly, mice were anesthetized and perfused with PBS through the heart, followed by 4% PFA in 0.1M PBS at pH 7.4. Interscapular BAT and epidydimal WAT were dissected and fixed overnight at 4°C. The samples were stored in 0.1% PFA at 4°C until processing. Fixed samples were dehydrated and embedded in paraffin. De-waxed sections were stained with an anti-tyrosine hydroxylase antibody (Cat#AB1542, Millipore Sigma, Burlington, MA), which was detected using the ABC method (Vector Laboratories, Newark, CA). Morphometric analyses were performed to assess the size of the lipid droplets. For this purpose, tissue sections were observed using a Nikon Eclipse E800 light microscope and digital images were acquired at 40X magnification with a Nikon DXM 1220 camera. For each sample, 5 tissue sections, at least 500 mm apart from each other, were selected. The size of at least 2,500 lipid droplets was measured for each sample (5 different areas per sample, 4 samples per condition). Data were analyzed using ImageJ.

#### Electron microscopy

Small pieces of WAT and BAT tissue were fixed in 2% glutaraldehyde, 2% paraformaldehyde in 0.1 M phosphate buffer, pH 7.4, overnight at room temperature. Samples were post-fixed in 1% osmium tetroxide, dehydrated, and embedded in epoxy resin. Thin sections were prepared with an MTX ultramicrotome (RMC, Tucson, AZ), stained with lead citrate, and imaged using a Philips CM10 transmission electron microscope (Philips, Eindhoven, Netherlands) as previously described (Colleluori et al., 2022).

### Quantitative real time reverse transcription PCR (RT-PCR)

First-strand cDNA was amplified using the TaqMan™ Universal PCR Master Mix, following manufacturer’s instructions. Primers and fluorogenic probes were purchased from Applied Biosystems (TaqMan(R) Gene Expression Assays, Foster City, CA) (Method Table M2) and performed in 96-well plates using a CFX96™ Real-Time System (Bio-Rad, Hercules, CA). Thermal cycling conditions were as follows: initial denaturation step at 95°C for 10 min, followed by 45 cycles, where each cycle was performed at 95°C for 30 s followed by 65°C for 60 s. Comparative quantitation of gene expression was conducted using the 2-ΔΔCt method, with vehicle control as the calibrator. Expression of target genes was normalized using the Best- keeper software (Pfaffl et al., 2004) using *Actb*, *Hprt*, and *Gapdh* (Schmittgen et al., 2008). Data are reported as fold change relative to control groups.

### Western blots

Western blot analyses were performed as described (Palese et al., 2019) with minor modifications. Briefly, proteins (30 µg) were denatured in SDS (8%) and β-mercaptoethanol (5%) at 95 °C for 5 min. After separation by SDS-PAGE on a 4–15% gel, the proteins were electrotransferred to nitrocellulose membranes. The membranes were blocked with 0.2% Tropix I-Block (Thermo Fisher Scientific) in PBS, pH 7.4, containing 0.1% Tween-20 at room temperature for 1 h. They were then incubated overnight at 4°C with either an anti-CB_1_ receptor rabbit monoclonal antibody (D5N5C, #93815, Cell Signaling Technology, Danvers, MA) or an anti-GAPDH rabbit monoclonal antibody (#ab181602, Abcam, Cambridge, UK) at 1:1,000 dilution in 0.2% Tropix I-Block in PBS, pH 7.4, containing 0.1% Tween-20, as the loading control. This was followed by incubation with a secondary horseradish peroxidase-linked antibody (1:5,000, Millipore Sigma) in Tris-buffered saline (TBS), pH 7.4, containing 0.1% Tween-20 at room temperature for 1h. Finally, proteins were visualized using an ECL kit (Bio-Rad, USA) and the chemiluminescence image was recorded using a LAS-4000 lumino-image analyzer system (Fujiflm, Tokyo, Japan).

### Transcriptomic analyses

*RNA isolation:* Total RNA was extracted as described (Lee et al, 2022). Samples with RNA integrity number ≥ 8.5 were used for RNA sequencing.

*RNA sequencing and bioinformatics analyses:* RNA sequencing was conducted at Novogene (Bejing, People’s Republic of China) using the Illumina NovaSeq platform with paired-end 150 bp (PE 150) sequencing strategy. Downstream bioinformatic analyses were performed using a combination of programs including STAR, HTseq, Cufflink and Novogene’s wrapped scripts, and alignments were parsed using STAR. Principal component analysis and comparative analyses of differentially expressed genes (DEGs) between test groups were performed using the DESeq2/edgeR package and a model based on negative binomial distribution. Resulting P values were adjusted using the Benjamini and Hochberg’s approach for controlling false discovery rate (Adjusted P values, Padj). For transcriptome analysis, comparative analysis of DEGs was carried out between two test groups. Changes displaying Padj < 0.05 were considered significant. DEG distribution was assessed using Volcano plots showing statistical significance (Padj) vs magnitude of change (fold change). DEGs were annotated using the Database for Annotation, Visualization and Integrated Discovery (DAVID) database, PANTHER gene ontology (GO) knowledgebase, and the Kyoto Encyclopedia of Genes and Genomes (KEGG) pathway database, which was implemented using the ClusterProfiler and/or iDep.93 bioinformatics platform. GO terms with adjusted P value less than 0.05 were considered significantly enriched in DEGs.

### Proteomic analyses

#### Tissue processing

WAT and BAT samples (*∼*50 mg) were transferred in ice-cold 2.0 ml Precellys soft tissue tubes in radioimmunoprecipitation lysis buffer (Cell Signaling Technology, Boston, MA) containing a protease inhibitor cocktail (Halt Protease, Thermo Fisher Scientific; 1/100 dilution) and a phosphatase inhibitor cocktail (Sigma Aldrich; 1/100 dilution). Samples were homogenized using a Bertin homogenizer at 4°C, 15 s/cycle for 2 cycles with 20 s pause between cycles and 45-s sample chilling time. The homogenates were transferred into 1.5 ml tubes, centrifuged for 15 min at 6,000 x *g* and 4°C, and the liquid collected under the fat layer was carefully transferred into another 1.5 ml tube using a 25G needle. The procedure was repeated twice. Protein concentration was measured using the bicinchoninic acid (BCA) method (Thermo Fisher Scientific), following manufacturer’s instructions.

#### Sample preparation for LC-MS/MS

The lysates were digested using a modified filter-assisted sample preparation protocol (Wiśniewski et al., 2009) over 10 kDa Microcon® centrifugal filters (Millipore Sigma). Samples were reduced on-filter using 4 mM TCEP (tris(2-carboxyethyl) phosphine) (Thermo Fisher Scientific) at room temperature for 30 min, followed by alkylation using 8 mM iodoacetamide (Sigma Aldrich) at room temperature for 30 min. The proteins were digested for 4 h at 37°C in 8 M urea using Lys-C (Wako Chemicals, Richmond, VA), followed by overnight trypsin digestion at 37°C after diluting the concentration of urea to <1.5 M. Peptide mixtures were cleaned using a Waters Sep-Pak tC18 cartridge, vacuum centrifuged, and resuspended in 3% acetonitrile/2% formic acid sample buffer prior to MS analysis. Peptide mixtures were subjected to LC-MS/MS analysis using an UltiMate 3000 RSLC (Thermo Fisher Scientific) coupled to an Orbitrap Fusion Lumos mass spectrometer (Thermo Fisher Scientific).

#### LC-MS/MS

LC separation was performed on a 50 cm×75 μm I.D. Acclaim® PepMap RSLC column. Peptides were eluted using a gradient of 3% to 22% acetonitrile in water containing 0.1% formic acid at a flow rate of 300 nL/min over 90 min. MS/MS spectra were extracted from RAW spectrometric files using PAVA (Guan et al., 2011) and were searched using Batch-Tag within Protein Prospector (v.6.3.5) against a decoy-containing database consisting of a normal *Mus musculus* Swissprot database concatenated with a randomized version (SwissProt.2019.4.8.random.concat, total of 17,016 protein entries). The mass accuracy for parent ions and fragment ions were set as ± 20 ppm and 0.6 Da, respectively. Trypsin was set as the enzyme, and a maximum of two missed cleavages were allowed. Protein *N*-terminal acetylation, methionine oxidation, and *N*-terminal conversion of glutamine to pyroglutamic acid were selected as variable modifications. The false detection rate for proteins and peptides was set at 1%. For each analysis, the number of unique peptides, total number of peptides, and protein coverage were determined. Quantitation of the proteins was performed in MaxQuant using similar settings as Protein Prospector searches. Briefly, RAW files were searched using MaxQuant (v. 1.6.0.16) against a FASTA containing the *Mus musculus* proteome obtained from the SwissProt open-source database (version December 2020). The first search peptide tolerance was set to 20 ppm, with main search peptide tolerance set to 4.5 ppm. The protein, peptide, and peptide spectrum match level false discovery rates were all 1%, as determined by a target-decoy approach. For quantification, intensities were determined as the full peak volume over the retention time profile. The degree of uniqueness required for peptides to be included in quantification was “Unique plus razor peptides.” The resulting label-free quantification (LFQ) values calculated using MaxQuant were used for comparing protein relative abundance among different samples.

### Metabolomic analyses

Interscapular BAT and epidydimal WAT were harvested and snap frozen in liquid N_2_ using a pre-cooled Wollenberger clamp (Wollenberger et al., 1960). Samples (*∼*50 mg) were pulverized to a homogeneous powder using a Cryomill (Retsch, Newtown, PA). An ice-cold mixture of methanol:acetonitrile:water (40:40:20, v/v; 0.5-0.6 mL) was added to ∼10-15 mg of powdered samples to make 25 mg/ml suspensions, which were centrifuged at 16,000 x *g* for 10 min at 4°C. Supernatants (3 µL) were analyzed as described (Jung et al., 2021). Briefly, a quadrupole-orbitrap mass spectrometer (Q Exactive Plus, Thermo Fisher Scientific) operated in negative ionization mode was coupled to a Vanquish Ultra High-Performance LC system (Thermo Fisher Scientific) with electrospray ionization. Scan range was *m/z* 70-1000, scanning frequency was 1 Hz and resolution was 140,000. LC separations were conducted using a XBridge BEH Amide column (2.1 mm x 150 mm, 2.5 mm particle size, 130Å pore size) with a gradient consisting of solvent A (20 mM ammonium acetate, 20 mM ammonium hydroxide in 95:5 water:acetonitrile, pH 9.45) and solvent B (acetonitrile). Flow rate was 150 µl/min. The gradient was: 0 min, 85% B; 2 min, 85% B; 3 min, 80% B; 5 min, 80% B; 6 min, 75% B; 7 min, 75% B; 8 min, 70% B; 9 min, 70% B; 10 min, 50% B; 12 min, 50% B; 13 min, 25% B; 16 min, 25% B; 18 min, 0% B; 23 min, 0% B; 24 min, 85% B; 30 min, 85% B. Autosampler temperature was 4°C. Data were analyzed using the MAVEN software (Melamud et al., 2010). To control for instrument variability, an internal standard, [^15^N]- valine, was spiked in the extraction solvent.

### Gut microbiota tests

#### Fecal Sample preparation

Mixed fecal droppings (∼2ml in volume) were collected from 4 mice in each cage at PND 44 (n=3 cages per group) and PND 70 (n=4 cages per group), Fecal samples were placed into 15 ml tubes and immediately stored in the -80 °C freezer. Samples were kept at -80 °C and thawed once to extract the DNA for 16S rRNA sequencing.

#### Preparation of DNA and 16S library construction for Illumina sequencing

Extraction of DNA from frozen stool samples was performed using the Qiamp DNA stool mini kit according to manufacturer’s instructions. Approximately 180-200 mg of stool sample was used for the DNA extraction. The resulting DNA was measured by Qubit and 5 ng was used as input for library construction. The library preparation was performed according to the Illumina 16S Metagenomic Sequencing Library Preparation protocol. More specifically, the protocol includes the primer pair sequences for the V3 and V4 region that create a single amplicon of approximately ∼460 bp [16S Amplicon Forward (V3 region): 5’-TCGTCGGCAGCGTCAGATGTGTATAAGAGACAGCCTACGGGNGGCWGCAG-3, and 16S Amplicon Reverse (V4 region): 5’- GTCTCGTGGGCTCGGAGATGTGTATAAGAGACAGGACTACHVGGGTATCTAATCC-3’]. The protocol also includes overhang adapter sequences that must be appended to the primer pair sequences for compatibility with Illumina index and sequencing adapters. Primers for the second step PCR reaction were used from the dual index kit for Nextera XT library construction. The resulting libraries were assayed for quantity using Qubit and for quality using the Agilent Bioanalyzer 2100 DNA HS chip. The libraries were normalized and then multiplexed together. The multiplexed library pool was quantified using qPCR and sequenced on Illumina Miseq 2X300bp run.

#### Analysis

We imported 3.4 million demultiplexed Illumina Miseq sequence reads, into QIIME2 version 2018.11 (https://qiime2.org; Bolyen et al., 2019). After quality control, we proceeded the analysis with the forward read only. We used DADA2 to denoise the single end forward reads with operational taxonomical units (OTUs), OTUs picked at 100% similarity. We assigned taxonomy to the OTUs with representatives’ sequences and the classifier trained with the q2-feature-classifier, classify-sklearn naïve Bayes taxonomy classifier against the greengenes database 13.8 99% OTUs reference sequences (Bokulich et al., 2018; McDonald et al., 2012). The QIIME2 created OTU table as well as the taxonomy table and metadata were transferred into R for statistical analysis (R version 4.0.2). We rarefied the OTU table via randomized sampling without replacement with 100 iterations at 124139 sequences per sample using the “EcolUtils” package (R core Team, 2018, https://www.r-project.org/; Salazar, G. 2020. EcolUtils: Utilities for community ecology analysis. https://github.com/GuillemSalazar/EcolUtils). We determined the effect of age and treatment and its interaction on microbial composition with Permutational multivariate analysis of variance (PERMANOVA) on a Bray Curtis dissimilarity matric that was generated from the rarefied OTU table using the adonis function of the vegan package version 2.5-6 in R. We performed a Shapiro-Wilk test to check for normality distribution of residuals for the Shannon diversity. Since the distribution was normal an ANOVA was used to check for significance of any of the factors for alpha diversity.

### Statistical analyses

Results are expressed as means ± SEM. Significance was determined using unpaired, two-tailed Student’s *t* test, or analysis of variance (ANOVA) (one way, two way) followed by Tukey’s or Bonferroni’s post hoc tests, as appropriate. GraphPad Prism version 8.0 (GraphPad Prism, San Diego, CA) was used to perform the analysis. Differences were considered significant if P < 0.05. Analysis of transcriptomics, metabolomics and proteomics results were conducted as described in previous sections.

### Tables for STAR Methods

**Method Table M1.** Acquisition parameters for Δ^9^-tetrahydrocannabinol (THC), anandamide (AEA), oleoylethanolamide (OEA), palmitoylethanolamide (PEA), 2-AG and their deuterium-containing analogs used as internal standards (ISTD).

| Compound | Precursor<br>(*m/z*) | Product<br>(*m/z*) | Dwell<br>time<br>(ms) | Fragmentat<br>ion (V) | Collision<br>(V) | ESI<br>Polarity |
| --- | --- | --- | --- | --- | --- | --- |
| THC | 315.25 | 193.1 | 200 | 145 | 21 | +ve |
| [<sup>2</sup>H<sub>3</sub>]-THC (ISTD) | 318.25 | 196.1 | 200 | 145 | 21 | +ve |
| OEA | 326.33 | 62.2 | 200 | 116 | 14 | +ve |
| [<sup>2</sup>H<sub>4</sub>]-OEA (ISTD) | 330.33 | 66.2 | 200 | 116 | 14 | +ve |
| PEA | 300.29 | 62.2 | 200 | 132 | 14 | +ve |
| [<sup>2</sup>H<sub>4</sub>]-PEA (ISTD) | 304.32 | 66.2 | 200 | 127 | 14 | +ve |
| AEA | 348.29 | 62.2 | 200 | 132 | 14 | +ve |
| [<sup>2</sup>H<sub>4</sub>]-AEA (ISTD) | 352.32 | 66.2 | 200 | 127 | 14 | +ve |
| 2-AG | 379.29 | 287.2 | 200 | 142 | 10 | +ve |
| [<sup>2</sup>H<sub>5</sub>]-2-AG (ISTD) | 384.33 | 287.2 | 200 | 142 | 10 | +ve |

**Method Table M2.**
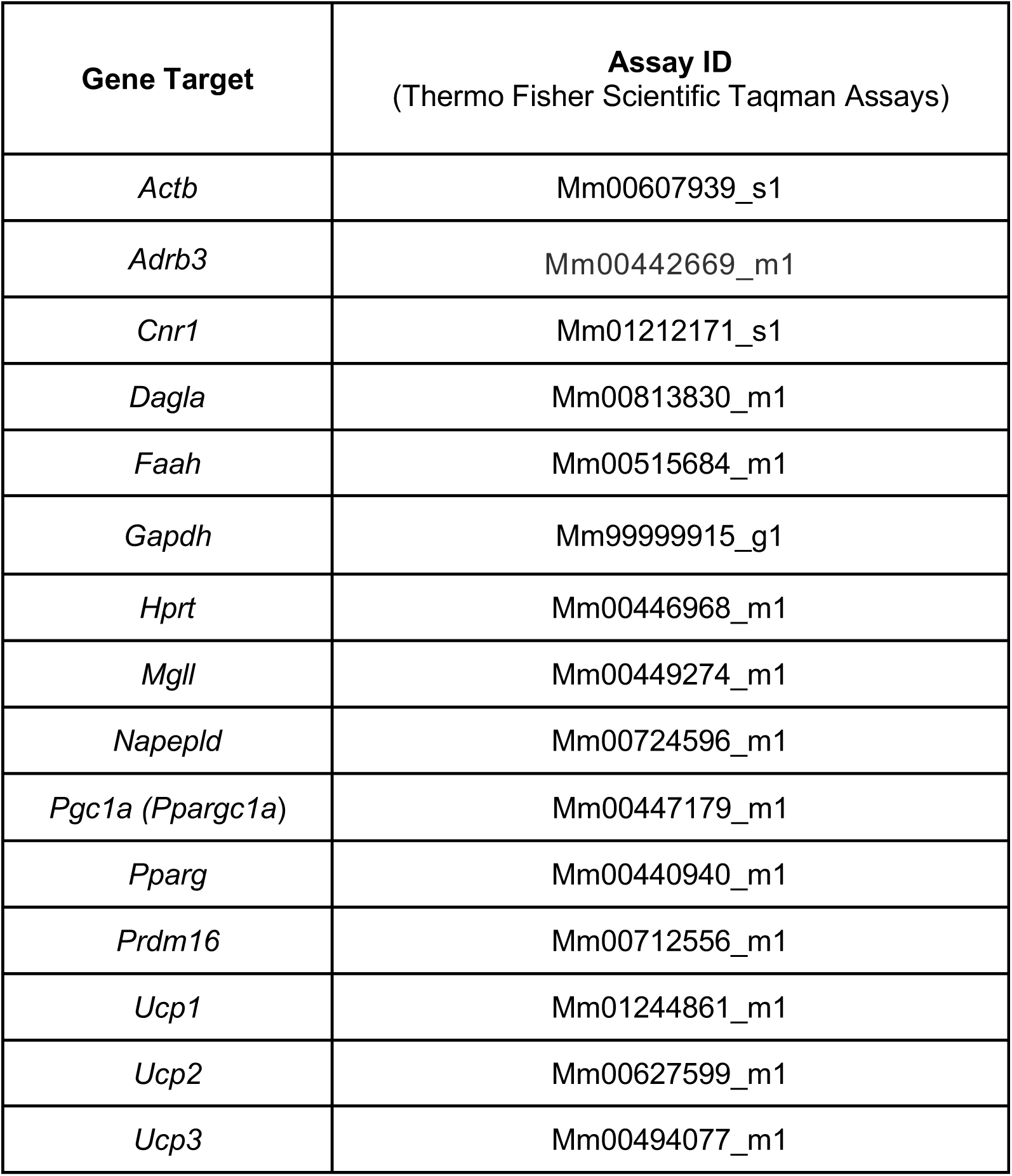
Quantitative RT-PCR probes used in the study.

## KEY RESOURCES TABLE

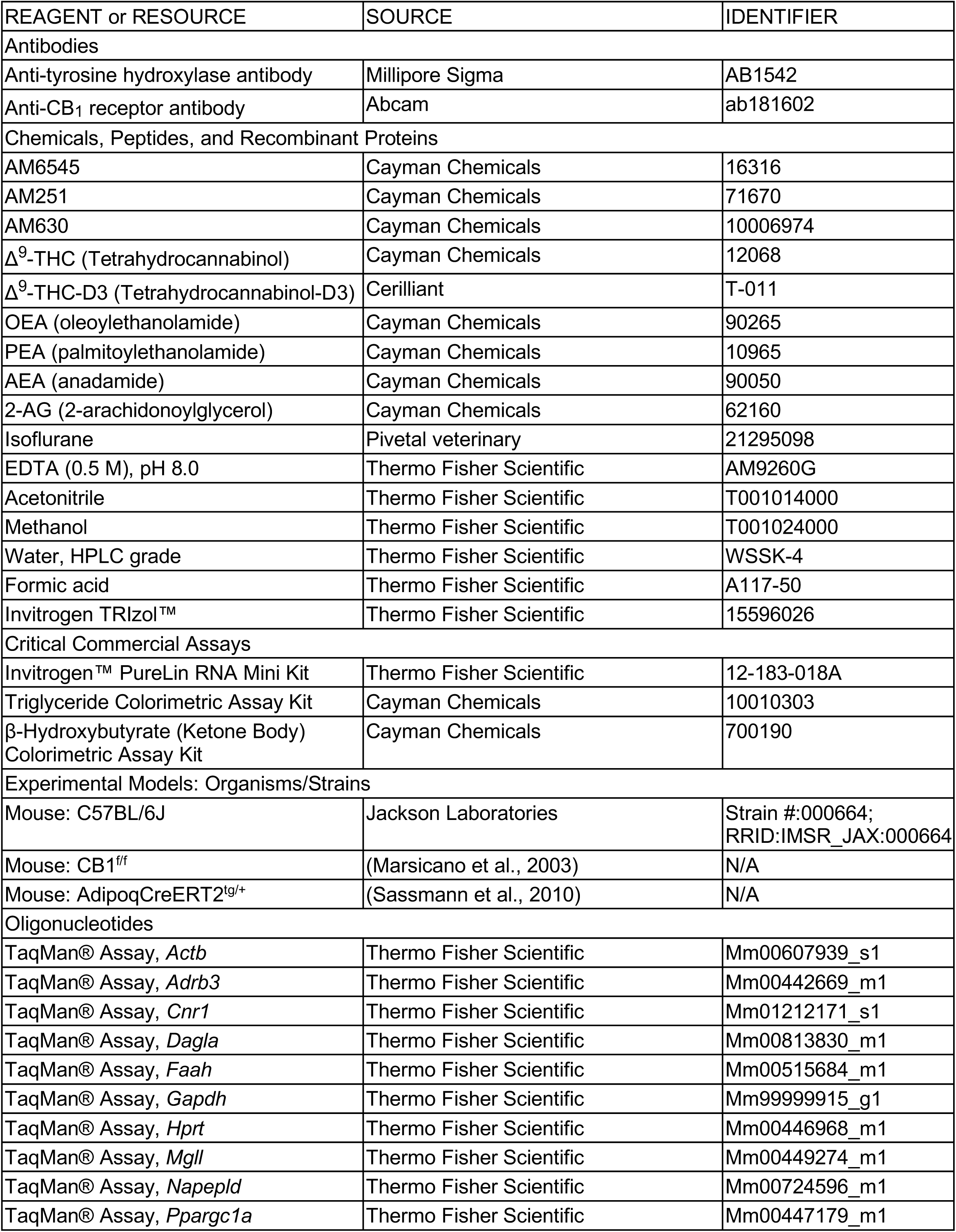
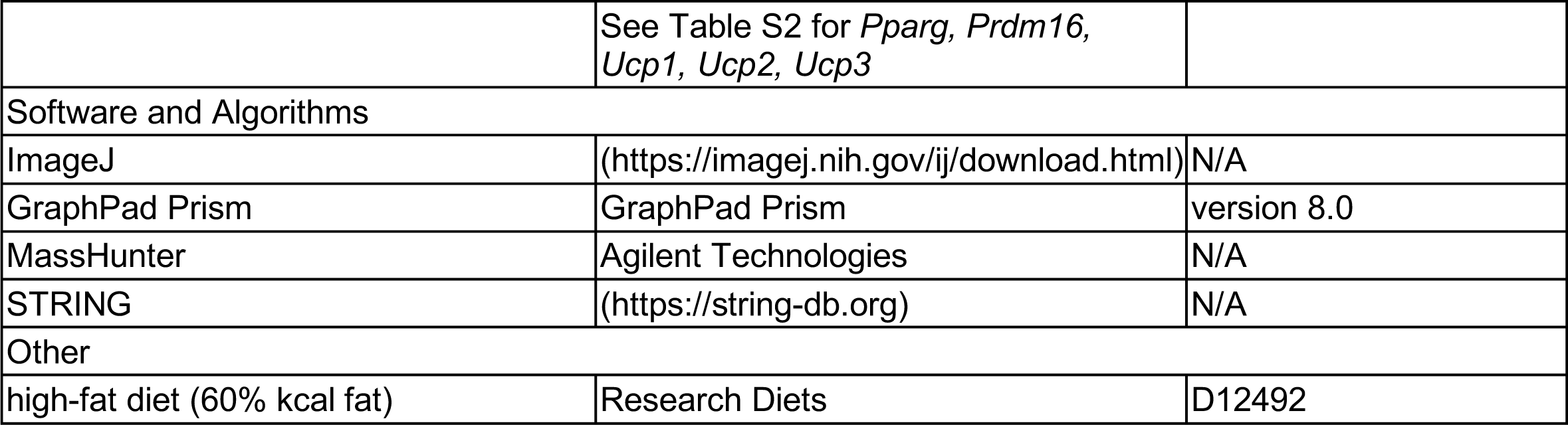

