## Supplemental Figures S1-S7, Supplemental Table S2,S3 for "ADOLESCENT EXPOSURE TO LOW-DOSE THC DISRUPTS ENERGY BALANCE AND ADIPOSE ORGAN HOMEOSTASIS IN ADULTHOOD"

Lin Lin et al.

#### List of Supplemental Information

- Supplemental Figures S1-S7
- Supplemental Tables S1-S7

**Supplemental Figure Legends**

**Supplemental Figure S1. Effects of adolescent THC exposure on body growth, motor activity, food intake, body water content, and microbiome composition.** Effects of adolescent THC or vehicle administration on **(A)** head length (n=4 per group); **(B)** tail length (n=4 per group); and **(C)** locomotor activity in a familiar environment (n=11 per group) in PND44 mice. No statistically significant differences were found by Student's *t* test (A, B) or two-way ANOVA followed by Bonferroni's post hoc test (C). **(D)** Circadian feeding behavior in PND44 mice (n=11 per group). Average data for 24 h are presented for **(D1)** Meal size; **(D2)** meal duration; **(D3)** meal frequency; **(D4)** latency to feed; **(D5)** post meal interval; and **(D6)** satiety ratio. The small change in satiety ratio (intermeal interval/meal size) did not affect overall food intake. *P < 0.05 by Student's *t* test. **(E)** Total water content (% body weight) on **(E1)** PND44 (n=16-17 per group); **(E2)** PND70 (n=15), and **(E3)** PND130, following 60 days of HFD (n=10-11). No statistically significant differences were found (Student's *t* test). **(F)** Adult intestinal microbiome composition (n=4 cages per group, each cage contained 4 mice). Statistical analyses are described under STAR Methods. Results are expressed as mean ± SEM.

**Supplemental Figure S2. Co-administration of $CB_2$ inverse agonist AM630 does not block the effect of THC on adolescent body weight gain. (A)** Body weight trajectory of adolescent male mice treated once daily with AM630 alone (1 mg/kg; gray circles) or THC (5 mg/kg) plus AM630 (green circles). **(B)** Cumulative body weight gain in the two groups. Results are expressed as mean ± SEM (n=7 per group). *P < 0.05, and **P < 0.01, two-way ANOVA followed by Bonferroni's post hoc test (A) or Student's *t* test (B).

**Supplemental Figure S3. Effects of adolescent THC exposure on the ECS in adipose organ.** Effects of adolescent THC or vehicle administration on critical ECS components in PND70 mice.

**(A)** $CB_1$ receptor mRNA (*Cnr1*) (RT-PCR, n=4 per group); **(B)** $CB_1$ receptor protein: left, representative Western blots; right, densitometry quantification (n=5). V, vehicle control; and T, THC-treated mice. **(C)** Diacylglycerol lipase-α mRNA (*Dagla*). **(D)** Monoglyceride lipase mRNA (*Mgll*). **(E)** N-acyl-phosphatidyl-ethanolamine phospholipase D mRNA (*Napepld*). **(F)** Fatty acid amide hydrolase mRNA (*Faah*) (n=3-4). Levels of endocannabinoid [anandamide (AEA), 2-AG] and endocannabinoid-like [palmitoylethanolamide (PEA), oleoylethanolamide (OEA)] substances in **(G)** BAT and **(H)** WAT (n=5-6). BAT, brown adipose tissue; WAT, white adipose tissue; AU, arbitrary units. Results are expressed as mean ± SEM. **P < 0.01, Student's *t* test.

**Supplemental Figure S4. Effects of adolescent THC exposure on select genes involved in adipose function.** RT-PCR analyses of BAT and WAT from PND70 mice treated in adolescence with vehicle (gray circles) or THC (blue squares). **(A)** *Pgc1a* (peroxisome proliferator-activated receptor-γ coactivator-1α). **(B)** *Prdm16* (PR/SET Domain 16). **(C)** *Adrb3* (β-adrenergic receptor). **(D)** *Ucp2* (Uncoupling protein 2). **(E)** *Ucp1* (Uncoupling protein 1). **(F)** *Ucp3* (Uncoupling protein 3). **(G)** *Pparg* (peroxisome proliferator-activated receptor-γ). BAT, brown adipose tissue; WAT, white adipose tissue; AU, arbitrary units. Results are expressed as mean ± SEM (n= 4 per group). *P < 0.05, and **P < 0.01, Student's *t* test.

**Supplemental Figure S5. Effects of adolescent THC exposure on BAT and WAT proteomes.** Untargeted proteomic analyses of adipose organ from PND70 mice treated in adolescence with vehicle. Volcano plots showing proteins that are upregulated (red, P<0.05), downregulated (green, P<0.05) or unchanged (black, P>0.05) in **(A)** BAT and **(B)** WAT. **(C)** Select proteins upregulated in BAT of THC- vs vehicle-treated mice (fold-change). Results are expressed as mean ± SEM (n=1-4). *P < 0.05, **P < 0.01, Student's *t* test.

**Supplemental Figure S6. Effects of adolescent THC exposure on intermediate metabolites in BAT and WAT. (A-D)** BAT: relative abundance of **(A)** glycolytic intermediates, **(B)** TCA cycle intermediates, **(C)** pentose phosphate pathway intermediates, and **(D)** nucleotides in PND70 mice treated with THC or vehicle during adolescence. **(E-K)** WAT: **(E)** principal component analysis of top 29 metabolites, and **(F-K)** relative abundance of indicated metabolites. Results are expressed as mean ± SEM (n=8). *P < 0.05, Student's *t* test.

**Supplemental Figure S7. Effects of adolescent THC exposure on adipocyte ultrastructure. (A)** Representative transmission electron microscopy images of interscapular brown adipocytes from THC-treated (left) and vehicle-treated (right) mice. Both samples show numerous mitochondria (some indicated by red arrows) that are well-organized, similar in shape and size, and with packed cristae (enlargement bottom left, some indicated with m), as typical of this adipose depot. **(B)** Lipid droplet size was similar in THC- and vehicle-treated mice. LD: lipid droplets; N: nucleus; Cap: capillary. Scale bar 5 μm. **(C)** Representative transmission electron microscopy image of mitochondria from epidydimal white adipocytes in vehicle-treated (upper panel) and THC-treated (middle and bottom panels) mice. LD: lipid droplets. Scale bar: 500 nm. **(D)** Density of noradrenergic fibers in BAT parenchyma. Light microscopy immunohistochemical staining of tyrosine hydroxylase-positive fibers (some indicated by red arrows) in interscapular BAT from THC-treated (left) and vehicle-treated (right) mice. Scale bar 30 μm.

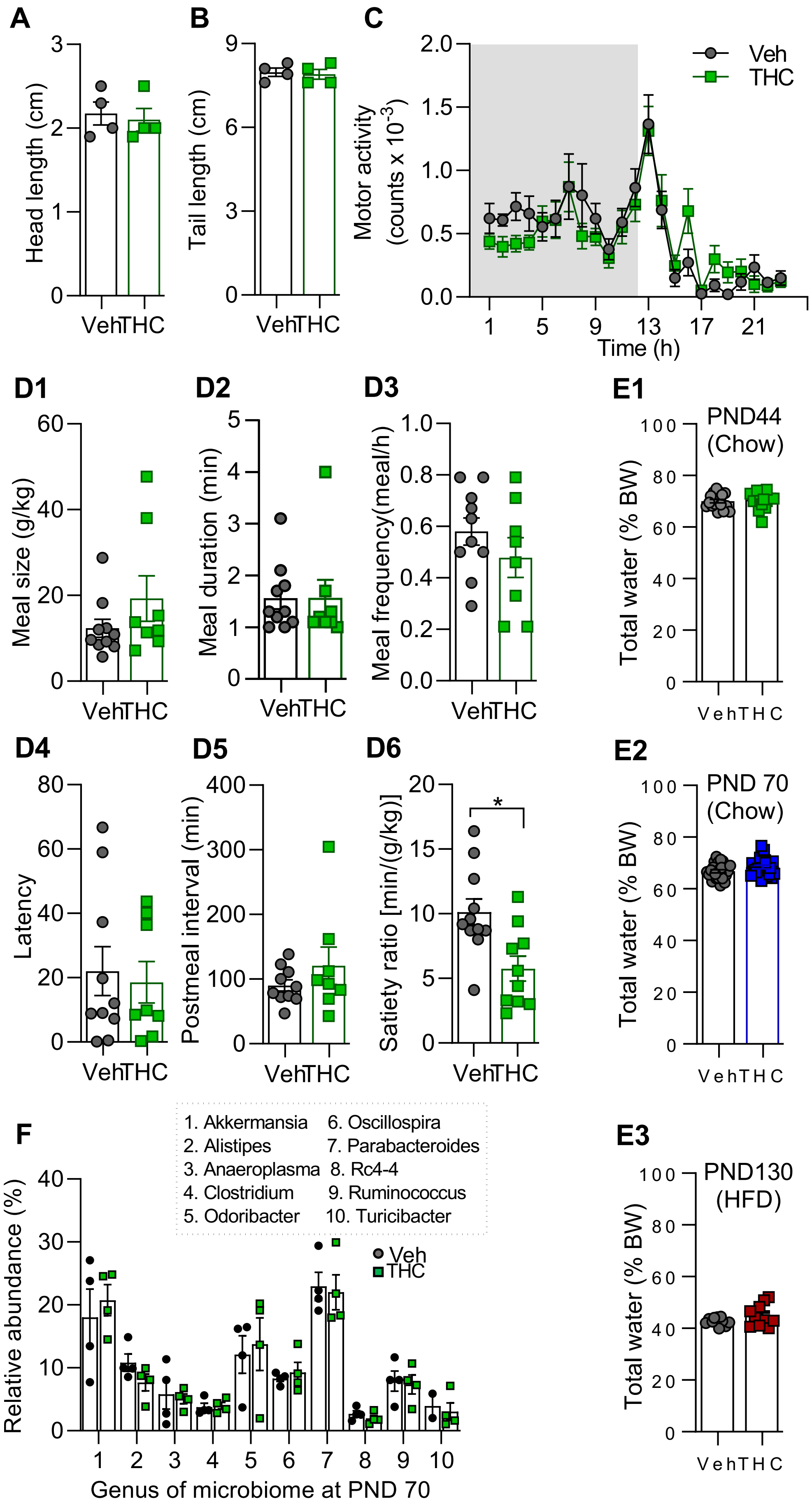


### Supplemental Figure S1

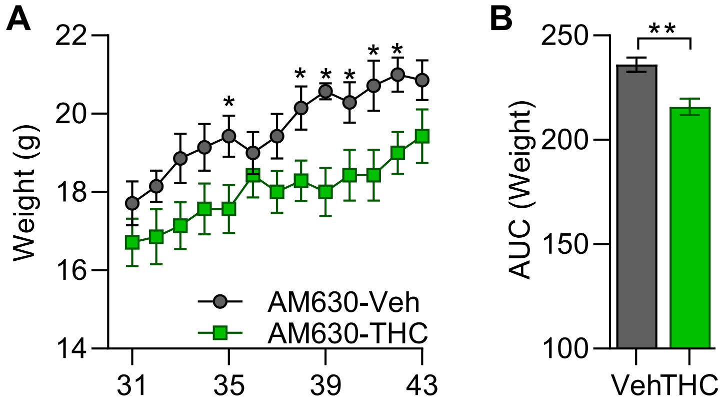


**Supplemental Figure S2**

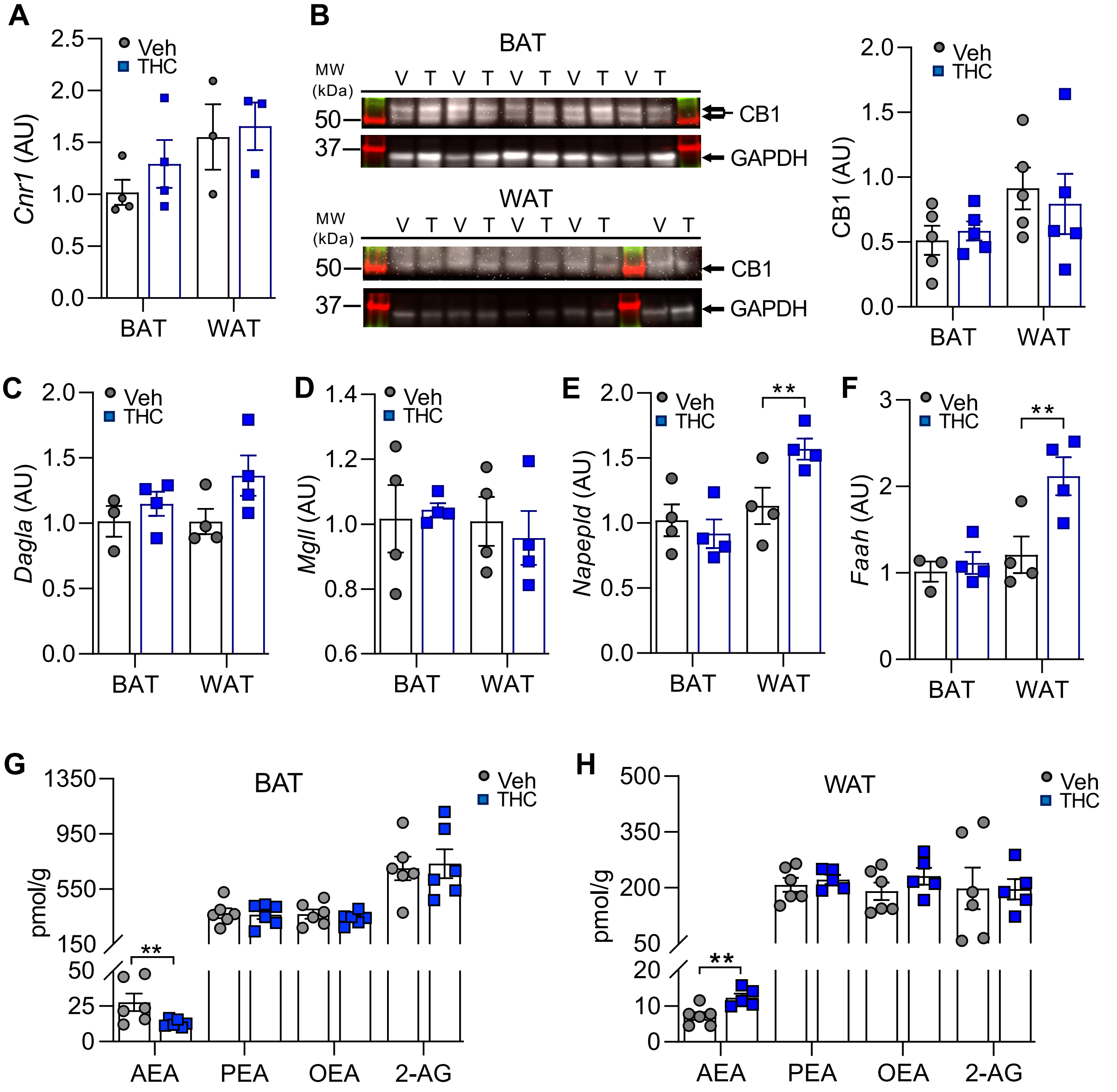


**Supplemental Figure S3**

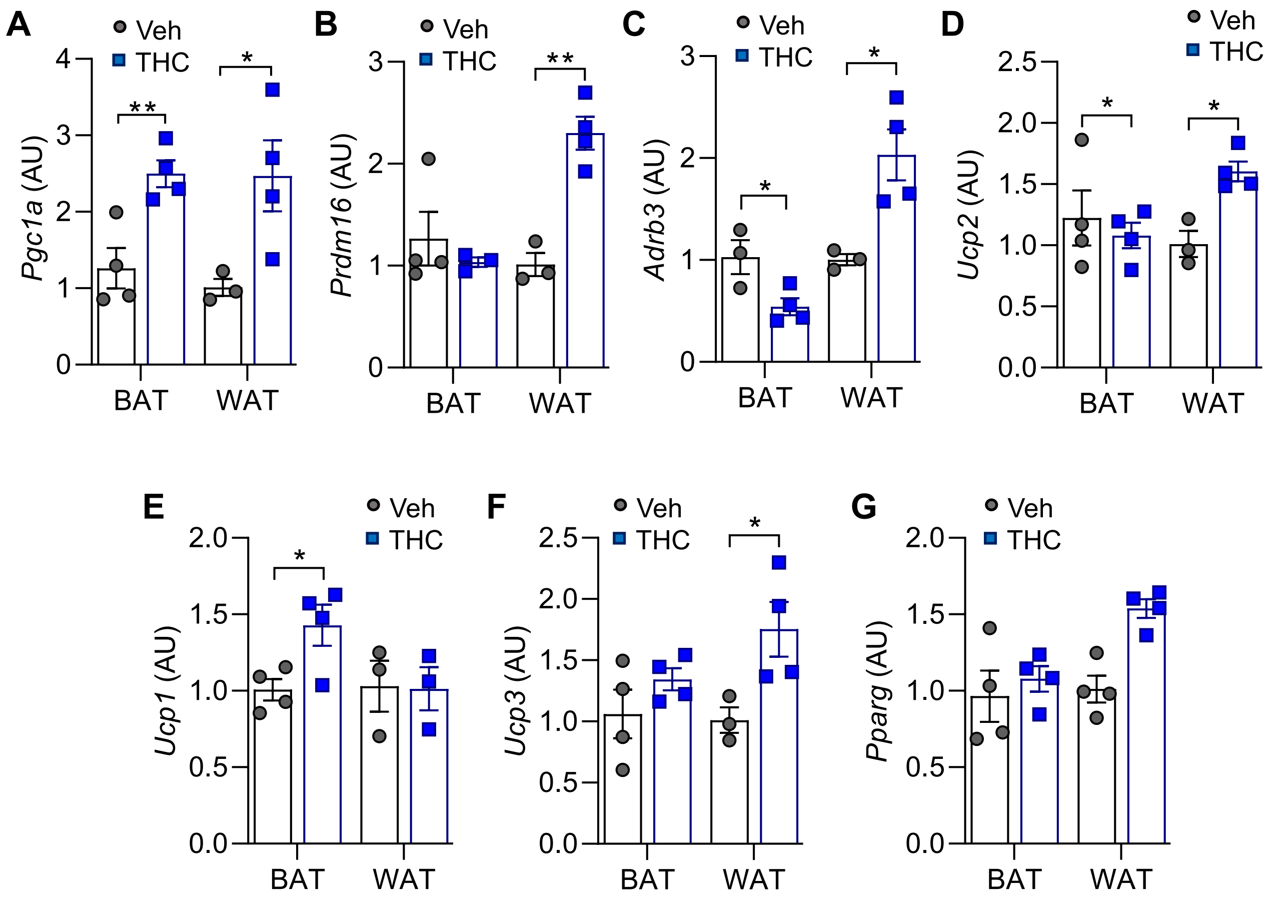


**Supplemental Figure S4**

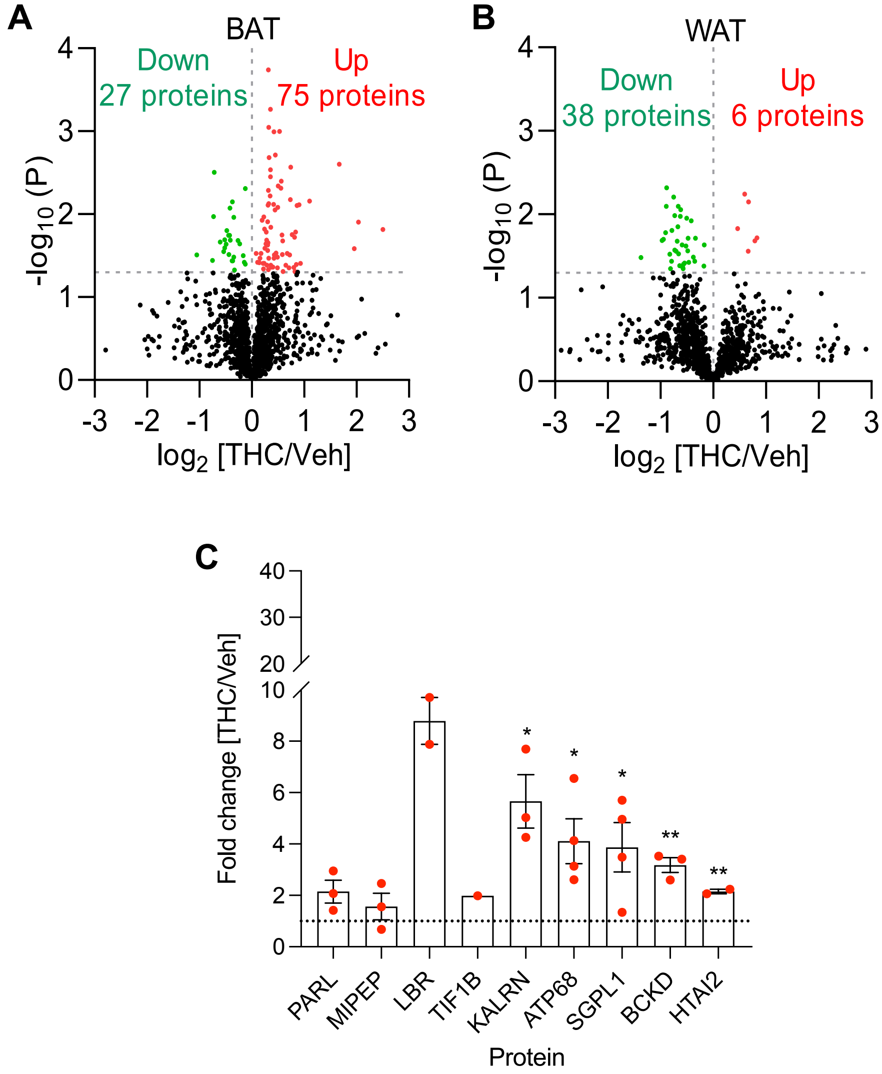


**Supplemental Figure S5**

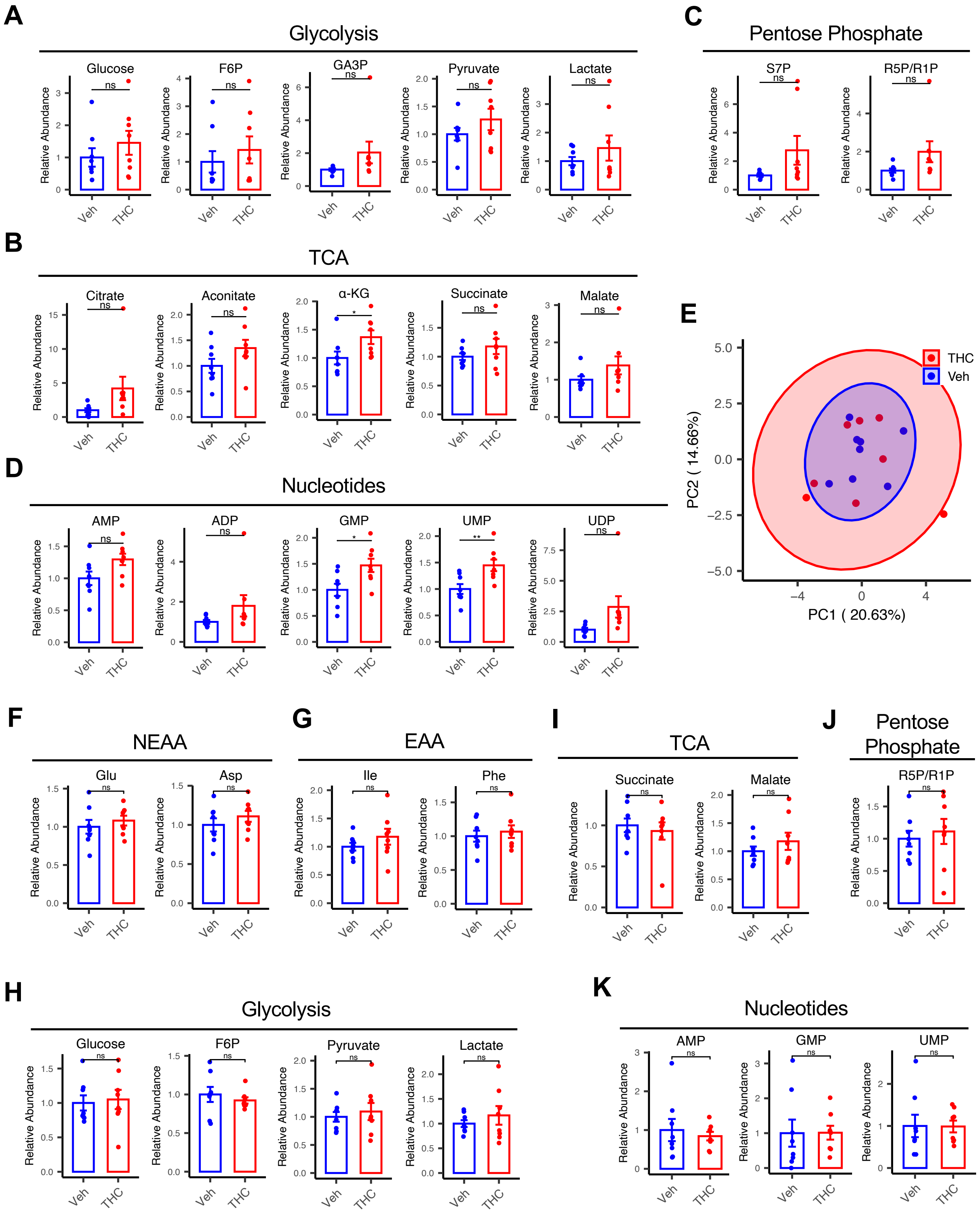


**Supplemental Figure S6**

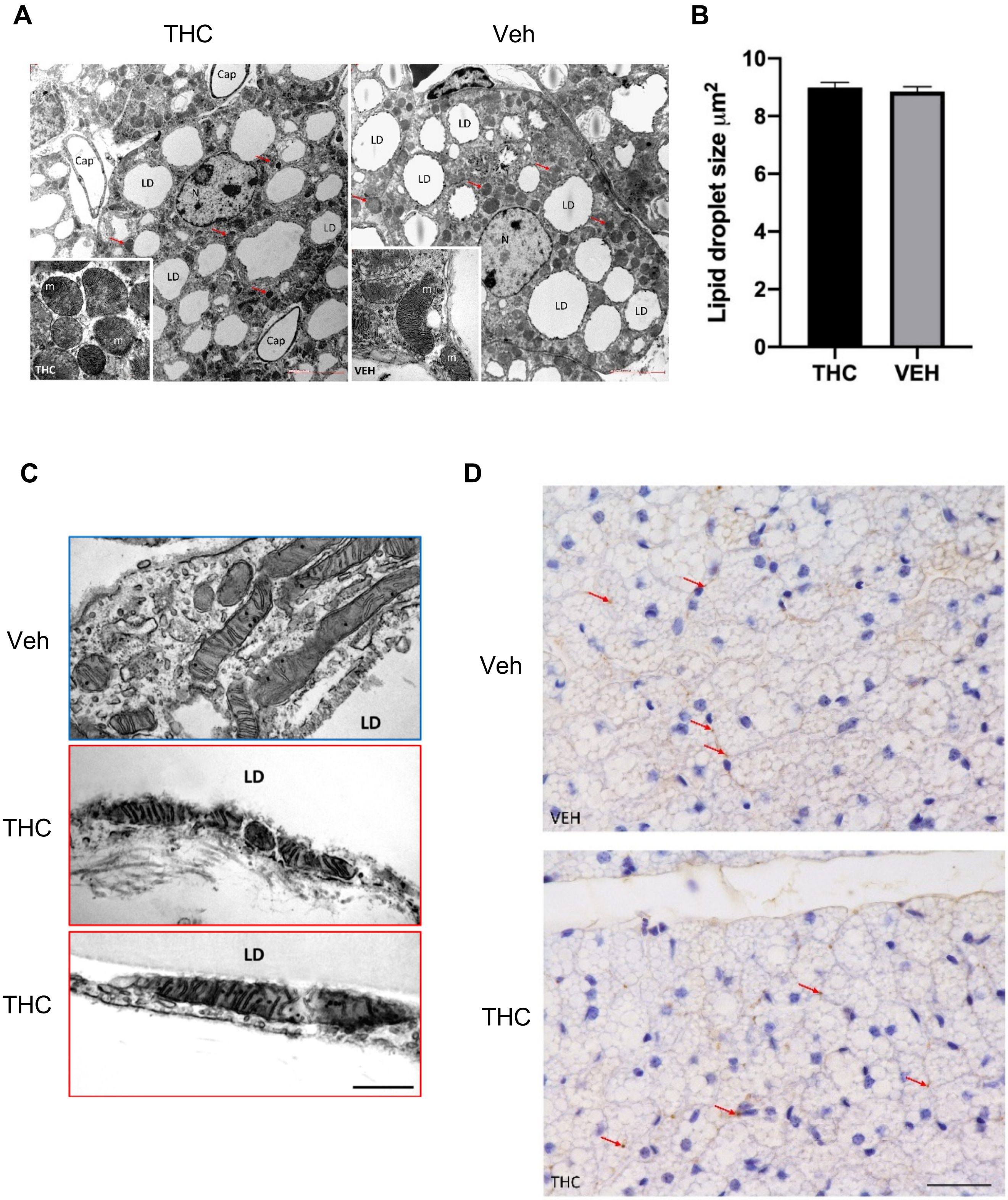


**Supplemental Figure S7**

#### Supplemental Tables

**Supplemental Table S1. Effects of adolescent THC exposure on adult intestinal microbiome composition** (Excel file containing dataset: Table_S1.xlsx)

**Supplemental Table S2. Effects of adolescent THC exposure on adult blood chemistry profile**

**Supplemental Table S3. Summary of metabolic changes produced by adolescent THC exposure**

**Supplemental Table S4. Transcriptome analysis of genes up- or down-regulated in BAT, WAT, and skeletal muscle of THC- vs vehicle-treated mice** (Excel file containing dataset: Table_S4.xlsx)

**Supplemental Table S5. STRING analysis of proteins upregulated in BAT and WAT of THC- vs vehicle-treated mice** (Excel file containing dataset: Table_S5.xlsx)

**Supplemental Table S6. List of proteins exclusively detected in BAT and WAT of THC-treated but not control mice** (Excel file containing dataset: Table_S6.xlsx)

**Supplemental Table S7. STRING analysis of proteins downregulated in BAT and WAT of THC- vs vehicle-treated mice** (Excel file containing dataset: Table_S7.xlsx)

**Supplemental Table S2. Effects of adolescent THC exposure on adult intestinal microbiome composition**

| **Fasted plasma** (Mean ± SEM) | **Vehicle** (n=5) | **THC** (n=5) | **P value** by student's *t* test |
|---|---|---|---|
| Total Protein (g/dL) | 4.1 ± 0.4 | 4.0 ± 0.4 | 0.8260 |
| Albumin (g/dL) | 2.3 ± 0.17 | 2.2 ± 0.2 | 0.5537 |
| Globulin (g/dL) | 1.7 ± 0.2 | 1.78 ±0.2 | 0.8904 |
| A/G Ratio | 1.4 ± 0.10 | 1.24 ± 0.05 | 0.2544 |
| AST (SGOT) ( IU/L) | 150.0 ± 8.1 | 114.8 ± 13.2 | 0.0876 |
| ALT (SGPT) ( IU/L) | 33.8 ± 5.4 | 33.8 ± 4.0 | 1.0000 |
| GGT (IU/L) | 1 ± 0 | 1.4 ± 0.4 | 0.3466 |
| Total Bilirubin (mg/dL) | 0.1 ± 0 | 0.14 ± 0.04 | 0.3466 |
| BUN (mg/dL) | 20.4 ± 2.2 | 19.8 ± 0.7 | 0.8028 |
| Creatinine (mg/dL) | 0.5 ± 0 | 0.5 ± 0 | 1.0000 |
| BUN/Creatinine Ratio | 40.8 ± 4.41 | 39.6 ± 1.5 | 0.8028 |
| Phosphorus (mg/dL) | 4.7 ± 0.40 | 5.22 ± 0.3 | 0.3501 |
| Glucose (mg/dL) | 96.6 ± 6.18 | 115.8 ± 10.3 | 0.1494 |
| Magnesium (mEq/L) | 0.18 ± 0.05 | 0.22 ± 0.05 | 0.5796 |
| NA/K Ratio | 5.8 ± 1.5 | 4.6 ± 0.51 | 0.4697 |
| Chloride (mEq/L) | 86.4 ± 4.39 | 83.4 ± 4.3 | 0.6379 |
| Cholesterol (mg/dL) | 75 ± 7.99 | 76.2 ± 12.2 | 0.9366 |
| Triglyceride (mg/dL) | 69 ± 7.65 | 67.8 ± 6.61 | 0.9085 |
| Amylase (IU/L) | 582.8 ± 117.2 | 1078.5 ± 290.8 | 0.2071 |
| PrecisionPSL™ (IU/L) | 40.8 ± 24.34 | 97.8 ± 54.33 | 0.3664 |
| Total Thyroid (T4) | 3.7 ± 0.3 | 3.6 ± 0.2 | 0.7625 |
| β-OH (Ketone body) (mM) | 1.02 ± 0.15 | 1.0 ± 0.12 | 0.939 |

**Supplemental Table S3. Summary of metabolic changes produced by adolescent THC exposure**

| **Parameter** | Ado-THC mice (**PND 44**, Chow-fed) | Ado-THC mice (**PND 70**, Chow-fed) | Ado-THC mice (PND 130, **HFD-fed**) |
|---|---|---|---|
| Food intake | ↔ | ↔ | ↔ |
| Motor activity | ↔ | ↔ | ↔ |
| Absorption | ↔ | ↔ | ↔ |
| Body weight | ↓ | ↔ | ↓ |
| Fat mass | ↔ | ↓ | ↓ |
| Lean mass | ↔ | ↑ | ↑ |
| EE | ↑ (Day & Night) | ↓ | ↑ (Night) |
| RER | ↓ (Day & Night) | ↓ | ↓ (Day) |
